# Modelling the viral dynamics of the SARS-CoV-2 Delta and Omicron variants in different cell types

**DOI:** 10.1101/2023.03.15.529513

**Authors:** Clare P. McCormack, Ada W. C. Yan, Jonathan C. Brown, Ksenia Sukhova, Thomas P. Peacock, Wendy S. Barclay, Ilaria Dorigatti

## Abstract

We use viral kinetic models fitted to viral load data from *in vitro* studies to explain why the SARS-CoV-2 Omicron variant replicates faster than the Delta variant in nasal cells, but slower than Delta in lung cells, which could explain Omicron’s higher transmission potential and lower severity. We find that in both nasal and lung cells, viral infectivity is higher for Omicron but the virus production rate is higher for Delta. However, the differences are unequal between cell types, and ultimately leads to the basic reproduction number and growth rate being higher for Omicron in nasal cells, and higher for Delta in lung cells. In nasal cells, Omicron alone can enter via a TMPRSS2-independent pathway, but it is primarily increased efficiency of TMPRSS2-dependent entry which accounts for Omicron’s increased activity. This work paves the way for using within-host mathematical models to understand the transmission potential and severity of future variants.

## Introduction

Since its designation as a variant of concern (VOC) by the World Health Organization on 26^th^ November 2021^1^, the Omicron (B.1.1.529\BA.1) variant of severe acute respiratory coronavirus 2 (SARS-CoV-2) has rapidly displaced the Delta (B.1.617.2) variant to become the dominant SARS-CoV-2 variant globally^2–4^. Analyses have demonstrated that Omicron can partially evade the immunity generated through previous infection and vaccination^5–8^, thereby leading to reduced vaccine effectiveness against symptomatic disease^9^.

Despite this reduction in vaccine effectiveness against symptomatic disease, the risk of severe outcomes (including hospital admission and death) following infection with Omicron is substantially lower than following infection with Delta, for both vaccinated and unvaccinated individuals^10,11^. While much remains to be understood about the mechanisms underpinning this observation, a reduction in the capacity of Omicron relative to Delta to replicate in lung cells has been suggested as a possible explanation^12^. On the other hand, Omicron’s increased ability to replicate in nasal cells has been suggested as an explanation for Omicron’s transmission advantage over Delta observed from epidemiological data^13–15^.

It has been hypothesised that these differences in viral replication capacity in different cells can be attributed to how Omicron and Delta utilise different pathways to enter host cells^12,16–18^. All SARS-CoV-2 viruses can enter cells which express both ACE2 and TMPRSS2 proteins, via fusion of the viral and cell membranes or via early endosomes^19^. However, Omicron is also able to efficiently enter ACE2+ cells via fusion from the endosome after endocytosis, without the involvement of TMPRSS2^16,20^. This could expand the range of cells that Omicron can infect. A complication is by entering cells via the endosome, SARS-CoV-2 may be inhibited by endosomal restriction factors such as interferon-induced IFITM proteins^21,22^ or NCOA7^23^. Understanding how virological properties shape the viral dynamics of SARS-CoV-2 in different cell types can enhance our understanding of the mechanisms underlying the observed differences in transmissibility between the Omicron and Delta variants. The different entry pathways and their inhibition by endosomal restriction factors are shown in Figure 1.

**Figure 1.**
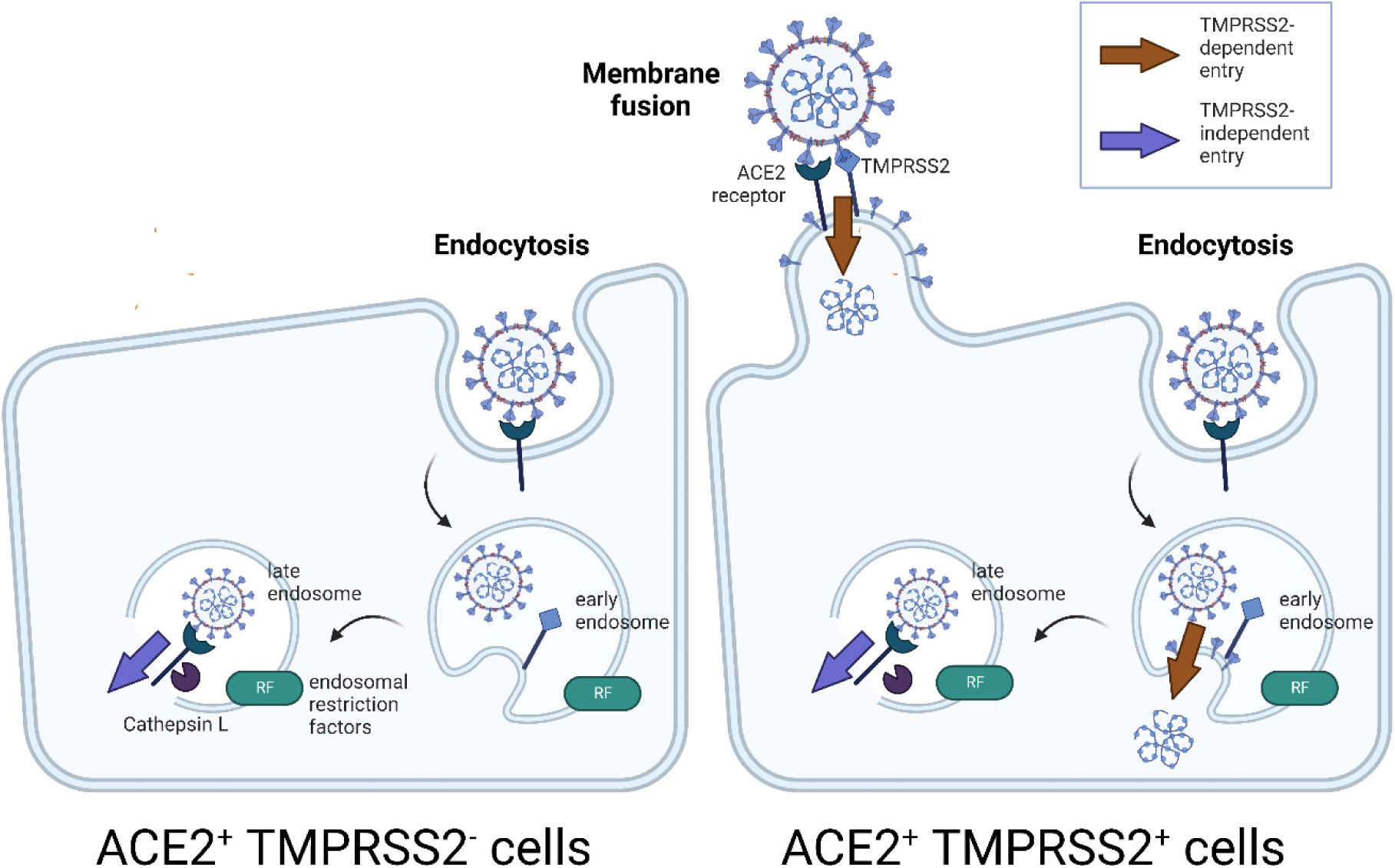
Illustration of TMPRSS2-dependent and TMPRSS2-independent entry pathways and their inhibition by endosomal restriction factors. In ACE2+ TMPRSS2+ cells (right), virus can enter in a TMPRSS2-dependent way via membrane fusion either at the cell surface or the early endosome; alternatively, virus can enter in a TMPRSS2-independent way through the late endosome. In ACE2+ TMPRSS2-cells (left), only the TMPRSS2-independent pathway is available. Both pathways can be inhibited by endosomal restriction factors. Created with BioRender.com.

Mechanistic mathematical models, calibrated against virological data, are a powerful tool for exploring viral dynamics as they provide a framework to quantify key characteristics of different variants and build an understanding of the drivers of inter-individual variation in response to infection. For example, previous within-host modelling studies of SARS-CoV-2 have enabled an understanding of the effect of covariates such as age, sex, and disease severity on viral load^24^, the degree of heterogeneity between individuals^25^, and the potential effect of antivirals and masking on viral load^26–30^. Models fitted to data from vaccine trials have suggested correlates of protection^31^, while more theoretical immunological models have also been developed with the ultimate aim of understanding the interplay between the immune response and disease severity^32–34^. Studies linking the within- and between-host scales have improved the understanding of the relationship between Ct values and infectiousness^25,35^, suggested characteristics of optimal testing regimes^36^ and provided explanations for population-level epidemiological observations, e.g. the apparent reduction in viral load observed during the declining phase of an epidemic^37^.

The studies mentioned above have all fitted their models to data from *in vivo* studies. While dynamics in humans are of ultimate interest, modelling viral kinetics and the immune response *in vivo* is extremely complex. Moreover, studies in humans are often limited, with little or no information on the timing and amount of the viral inoculum, sparse longitudinal observations, and often no available measurements before the start of symptom onset^25,35^. *In vitro* studies offer a simplification of *in vivo* dynamics, where both the inoculum and method of inoculation is controlled, and where some components of the immune response – such as the adaptive immune response – are eliminated. Viral load can be measured from the start of infection, which is important for estimating the growth rate and basic reproduction number of the virus, which in turn can be used to assess the potential for load variants to outcompete existing variants.

Here, using viral kinetics models calibrated against viral replication data generated through experimental infection studies in primary human nasal epithelial cells (hNECs) and immortalised Calu-3 lung cells, we characterise and compare the dynamics of Omicron and Delta viruses in each cell type by quantifying key properties such as the basic reproduction number and growth rate, and by exploring how these properties vary for different entry pathways, and in the presence and absence of functional endosomal restriction factors.

## Methods

### Experimental Design/Data

Full details of the viral kinetics experiments conducted are provided in Peacock et al.^20^. In brief, Calu-3 and primary human nasal epithelial cells (hNECs) were inoculated with Omicron BA.1 and Delta/B.1.617.2 isolates at an MOI of 0.001 (Calu-3) or 0.05 (hNECs), and incubated for 1 hr at 37°C. The inoculum was then removed and, in the case of Calu-3 cells, replaced with 1 mL serum-free DMEM. For Calu-3 cells, 100 μL of supernatant was collected for titration at 18, 24, 48 and 72 hours post-infection. For hNECs, at the same timepoints post-infection, the supernatant was collected by adding 200 μL of serum-free DMEM to the apical surface, incubating for 10 minutes, and removing for collection.

Additional experiments were performed to investigate the effect of Camostat mesylate (henceforth referred to as Camostat) and Amphotericin B on viral kinetics. Camostat is a serine protease inhibitor and thus inhibits TMPRSS2, while Amphotericin B inhibits the restriction of viral endosomal entry by endosomal restriction factors such as IFITM proteins^38–40^. Cells were pre-treated both basolaterally and apically with either 50 μM of Camostat, 1μM of Amphotericin B or no drug for 2 hours prior to infection, and this concentration of drug was maintained in the basolateral media throughout the experiment. Infections and collection of timepoints were performed as above.

All experiments were performed in triplicate. Infectious viral titres were quantified by plaque assay and viral genome numbers were quantified by E gene RT-qPCR.

### Mathematical Models

#### Model 1

To gain an initial understanding of the differences between the observed infection dynamics of the Omicron and Delta variants across each cell type, we first developed a simple model of the viral dynamics which assumed a single virus entry pathway for both variants. A simple schematic of the model is provided in Figure 2. Model equations are in the Supplementary Methods.

**Figure 2:**
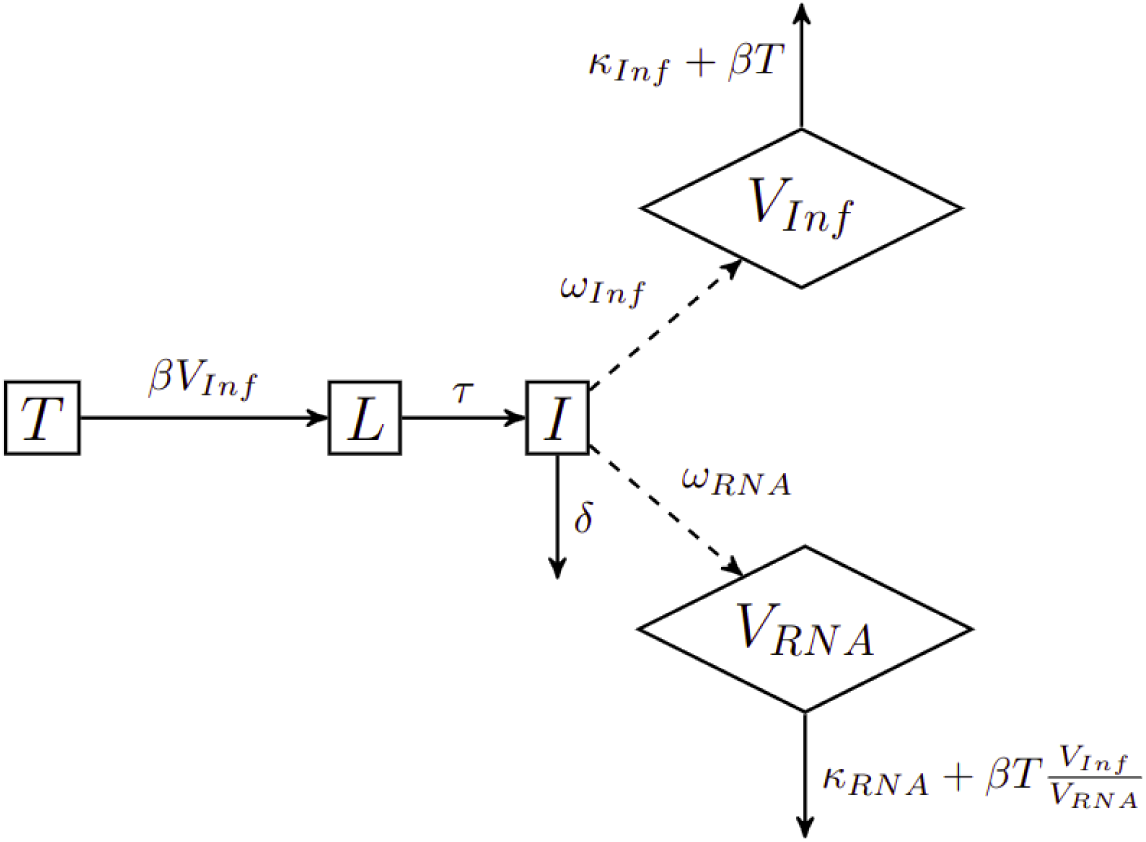
Compartmental diagram of model 1. Target cells *T* become infected at a rate *β* per infectious virion (V_Inf_). Following an eclipse phase of mean duration of 1/τ days, infectious cells (I) produce infectious virus as measured by plaque-assay (V_Inf_) and total virus as measured by qPCR (V_RNA_) at rates ω_Inf_ and ω_RNA_ respectively. Infected cells have a mean lifespan of 1/δ days, and infectious and total virus are assumed to decay at rates κ_Inf_ and κ_RNA_ respectively.

Target cells *T* become infected at a rate *β* (target cell infection rate) per infectious virion (V_Inf_). The target cell infection rate is defined as the proportion of cells infected per day, per unit inoculum of 1 pfu/mL. Following an eclipse phase of mean duration of 1/τ days, infectious cells (I) produce infectious and non-infectious virus. Infectious virus (V_Inf_), as measured by plaque-assay, and total (infectious and non-infectious) virus (V_RNA_), as measured by RT-qPCR, are produced at rates ω_Inf_ and ω_RNA_ respectively. (While the RT-qPCR assay could detect viral RNA released from lysed cells as well as from extracellular viral particles, the model assumes that all detected viral RNA originates from viral particles.) The duration of the eclipse phase 1/τ reflects the speed of a single viral replication cycle, whereas the rate of viral production *ω_Inf_* also reflects replication capacity. Infectious cells have a mean lifespan of 1/δ days, and infectious and non-infectious virus is assumed to decay at rates κ_Inf_ and κ_RNA_ respectively, with additional loss due to entry into cells. Cells which express the ACE2 receptor, which is required for SARS-CoV-2 virus entry, are considered target cells.

#### Model 2

The experimental data showed that while Delta can only enter ACE2^+^ cells by utilizing the TMPRSS2 protein, Omicron can enter these cells both through TMPRSS2-dependent and TMPRSS2-independent pathways. Thus, in Model 2 we modify Model 1 to account for the two possible cell entry pathways, where TMPRSS2-independent pathways can be used to infect all ACE2^+^ cells, and TMPRSS2-dependent pathways can only infect ACE2^+^ TMPRSS2^+^ cells. We assume that all ACE2^+^ cells become infected through TMPRSS2-independent pathways at a rate *β_E_* per infectious virion (V_Inf_) and that ACE2^+^ TMPRSS2^+^ target cells (T_+_) also become infected through TMPRSS2-dependent pathways at a rate *β_T_* per infectious virion. We assume that endosomal restriction factors decrease the infectivity through the TMPRSS2-independent pathways by a factor *f_E_*, and through TMPRSS2-dependent pathways by a factor *f_T_*. Cells infected through TMPRSS2-independent and TMPRSS2-dependent pathways (*L_E_* and *L_T_* respectively) undergo eclipse phases of mean duration 1/*τ_E_* and 1/*τ_T_* days respectively. Following the eclipse phase, infectious cells (I) behave as per Model 1. The flow diagram of Model 2 is described in Figure 3.

**Figure 3:**
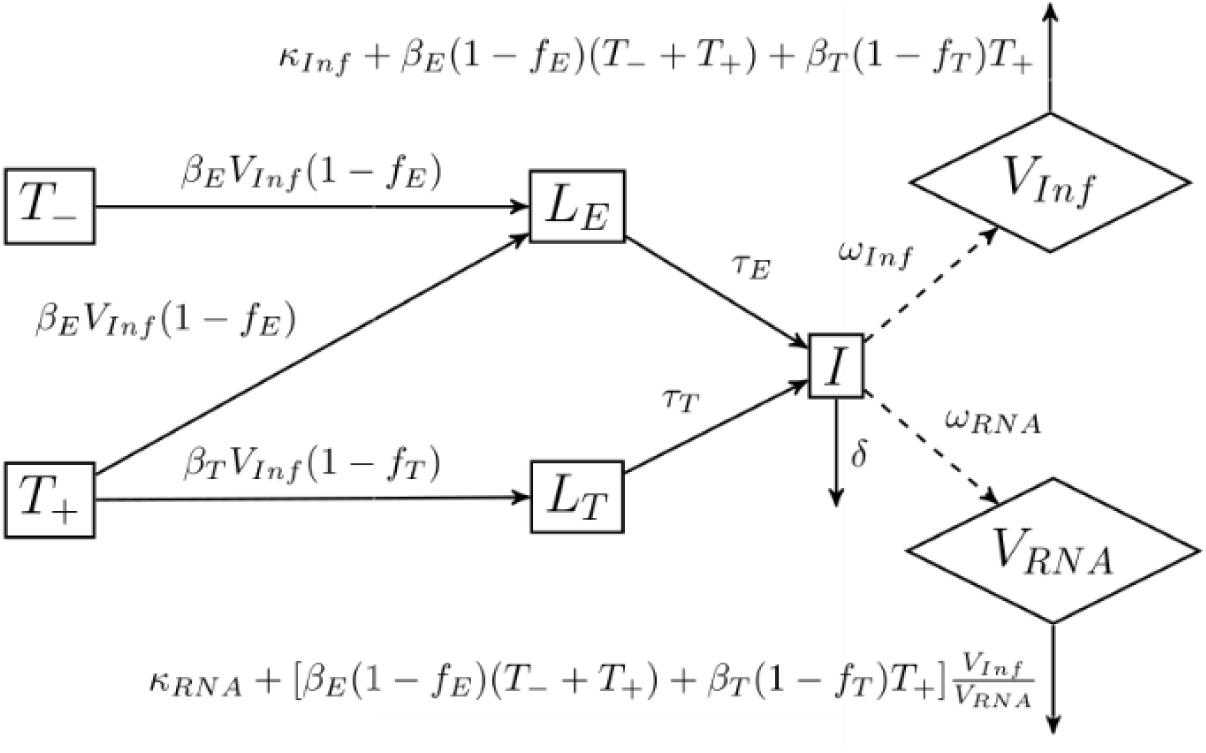
Compartmental diagram for Model 2. ACE2^+^ TMPRSS2^-^ target cells (T-) become infected through TMPRSS2-independent pathways at a rate *β_E_* per infectious virion (V_Inf_). ACE2^+^ TMPRSS2^+^ target cells (T+) become infected through TMPRSS2-independent pathways at a rate *β_E_* per infectious virion, and through TMPRSS2 -dependent pathways at a rate *β_τ_* per infectious virion. Endosomal restriction factors decrease the infectivity through TMPRSS2-independent pathways by a factor *f_E_*, and through TMPRSS2-dependent pathways by a factor *f_T_*. Cells infected through TMPRSS2-independent and TMPRSS2-dependent pathways (*L_E_* and *L_T_* respectively) undergo eclipse phases of mean duration *1/τ_E_* and *1/τ_T_* days respectively. Following the eclipse phase, infectious cells (I) behave as per Model 1.

Model 2 is fitted to hNEC and Calu-3 data without drug, in the presence of Camostat, in the presence of Amphotericin B, and in the presence of both drugs. For each virus strain/cell type combination, all data (with and without drugs) is fitted simultaneously. Camostat is assumed to completely inhibit TMPRSS2-dependent pathways (we set *β_T_ =* 0); although Camostat also inhibits other serine proteases, but as TMPRSS2 is the main serine protease involved in SARS-CoV-2 virus entry, and because the activation of S2’ in TMPRSS2-independent pathways likely occurs by cathepsin proteases rather than serine proteases, for simplicity we assume that Camostat does not affect TMPRSS2-independent pathways. Amphotericin B is assumed to completely remove endosomal restriction (we set *f_E_* = *f_T_* = 0).

#### Inferential framework

We calibrated the models to the observed data in a Bayesian inferential framework using Markov Chain Monte Carlo (MCMC) methods. In the results, we reported the median and 95% credible interval (CrI) of the parameter estimates. Full details of the model fitting algorithms are provided in the Supplementary Methods.

## Results

### Data Description

Figure 4 summarises the study data. In control wells with no drug, temporal trends in viral growth varied both between strains and cell type. In hNECs, Omicron grew more rapidly than Delta, with viral load peaking by 1 day post infection (p.i.) for Omicron, compared with 2 days p.i. for Delta. However, despite this initial growth advantage, viral load was higher for Delta compared to Omicron at 3 days p.i.. In Calu-3 cells, the peak in viral load was observed 2-3 days p.i. for both Omicron and Delta. In contrast with hNECs where the strain corresponding to the highest viral load was dependent on measurement time, infectious and non-infectious viral load was higher for Delta than Omicron across all time points for Calu-3 cells.

**Figure 4:**
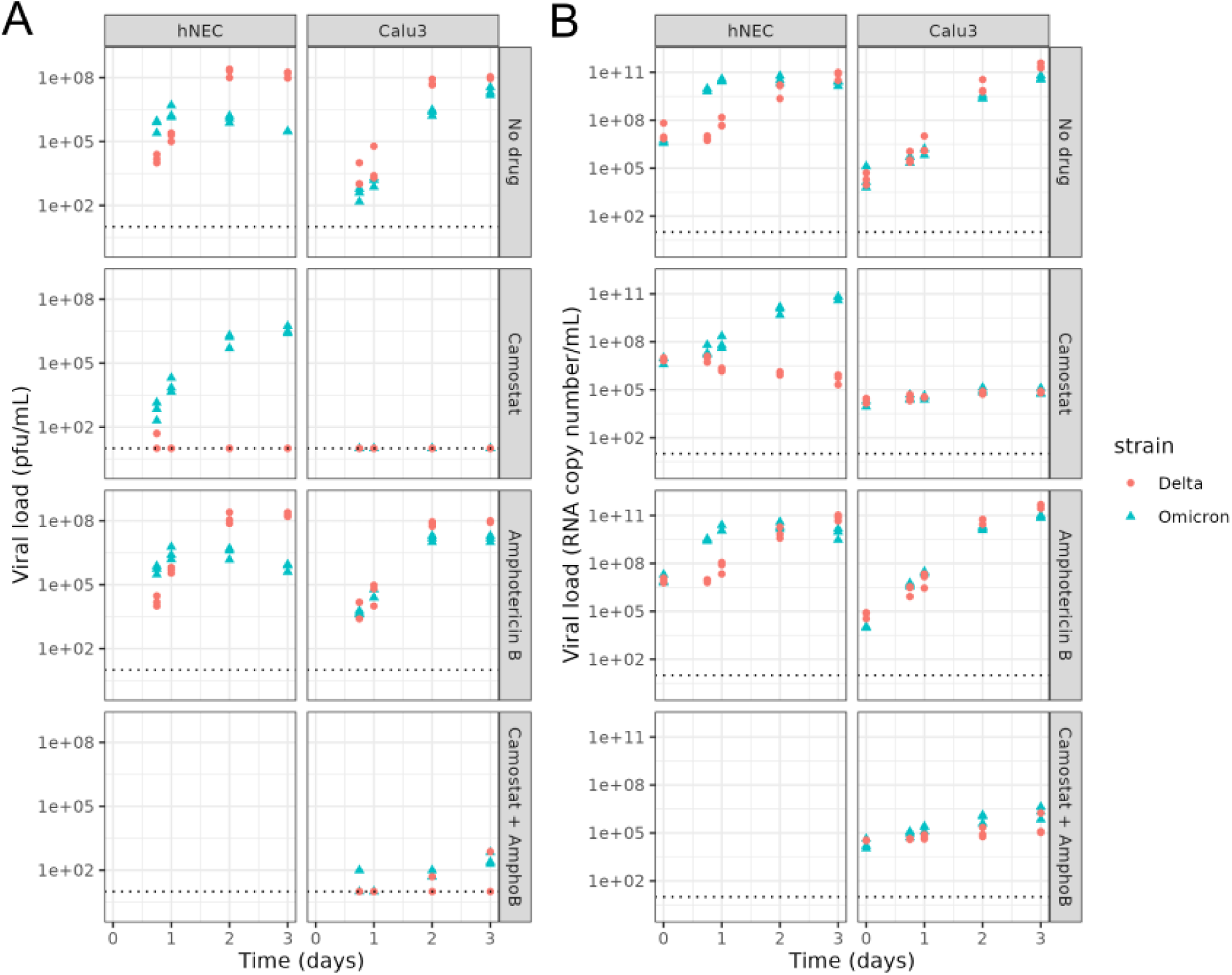
Summary of Data. (A) Infectious viral load (pfu/mL) quantified by plaque assay and (B) RNA copy number/mL quantified by RT-qPCR for Omicron (blue triangles) and Delta (red dots) for different cell types (columns) and drug treatments (rows). The dotted line shows the limit of detection; markers on the lines represent measurements below the limit of detection. Cells were infected on day 0.

In the presence of the drug Camostat which inhibits serine proteases, and thus TMPRSS2-dependent pathways, replication of Delta was severely inhibited in hNECs with no detectable increase in virus throughout the course of the experiment. In contrast, Omicron successfully replicated in hNECs in the presence of Camostat, albeit at a slower rate than in untreated wells. However, similar levels of viral load were obtained by 2-3 days p.i. in both the presence and absence of Camostat. For both strains, the presence of Amphotericin B had little impact on viral load dynamics in hNECs. In Calu-3 cells, infectious virus was not detected at any point for either Omicron or Delta in the presence of Camostat. In the presence of the drug Amphotericin B which is described as specifically inhibiting endosomal restriction^22,39^, Omicron infectious viral titres were consistently higher than in control wells, until 3 days p.i. when titres became similar. Delta geometric mean infectious viral titres were similar with and without Amphotericin B.

Experiments with both drugs combined were only performed for Calu-3 cells. For both virus strains, the infectious viral load was below the limit of detection in the presence of Camostat only. Adding Amphotericin B raised the infectious viral load to above the limit of detection for all replicates for Omicron, but only for some replicates for Delta (Figure 4).

### Viral Fitness

#### Model 1

For both Omicron and Delta, the simple model (Model 1) captured the viral dynamics observed in control wells of hNECs and Calu-3 cells (Figures 5, 6). For hNECs, we estimated a higher target cell infection rate *β* for Omicron relative to Delta (2.13e-05 ( 95% CrI: 1.54e-05, 3.07e-05) cell (pfu/mL)^-1^ day^-1^ for Omicron, 8.89e-08 (95% CrI: 5.91e-08, 1.28e-07) cell (pfu/mL)^-1^ day^-1^ for Delta), but a lower infectious virus production rate *ω_Inf_* for Omicron relative to Delta (4.07e+03 (95% CrI: 3.13e+03, 5.50e+03) pfu/mL cell^-1^ day^-1^ for Omicron, 3.42e+05 (95% CrI: 2.41e+05, 5.04e+04) pfu/mL cell^-1^ day^-1^ for Delta). All estimated parameters are presented in Supplementary Table 2. These results suggest an approximately 1.5-fold difference in the growth rate between these strains in hNECs, with a growth rate of 11.61 day^-1^ (95% CrI: 10.43,12.99) for Omicron and 6.87 day^-1^ (95% CrI: 6.33,7.50) for Delta. Furthermore, we estimated a higher basic reproduction number (*R*_0_) for Omicron (65.04 (95% CrI: 51.94, 82.71)) compared with Delta (23.13 (95% CrI: 19.91, 27.23)), thereby suggesting an overall increase in viral fitness of Omicron relative to Delta in hNECs.

**Figure 5:**
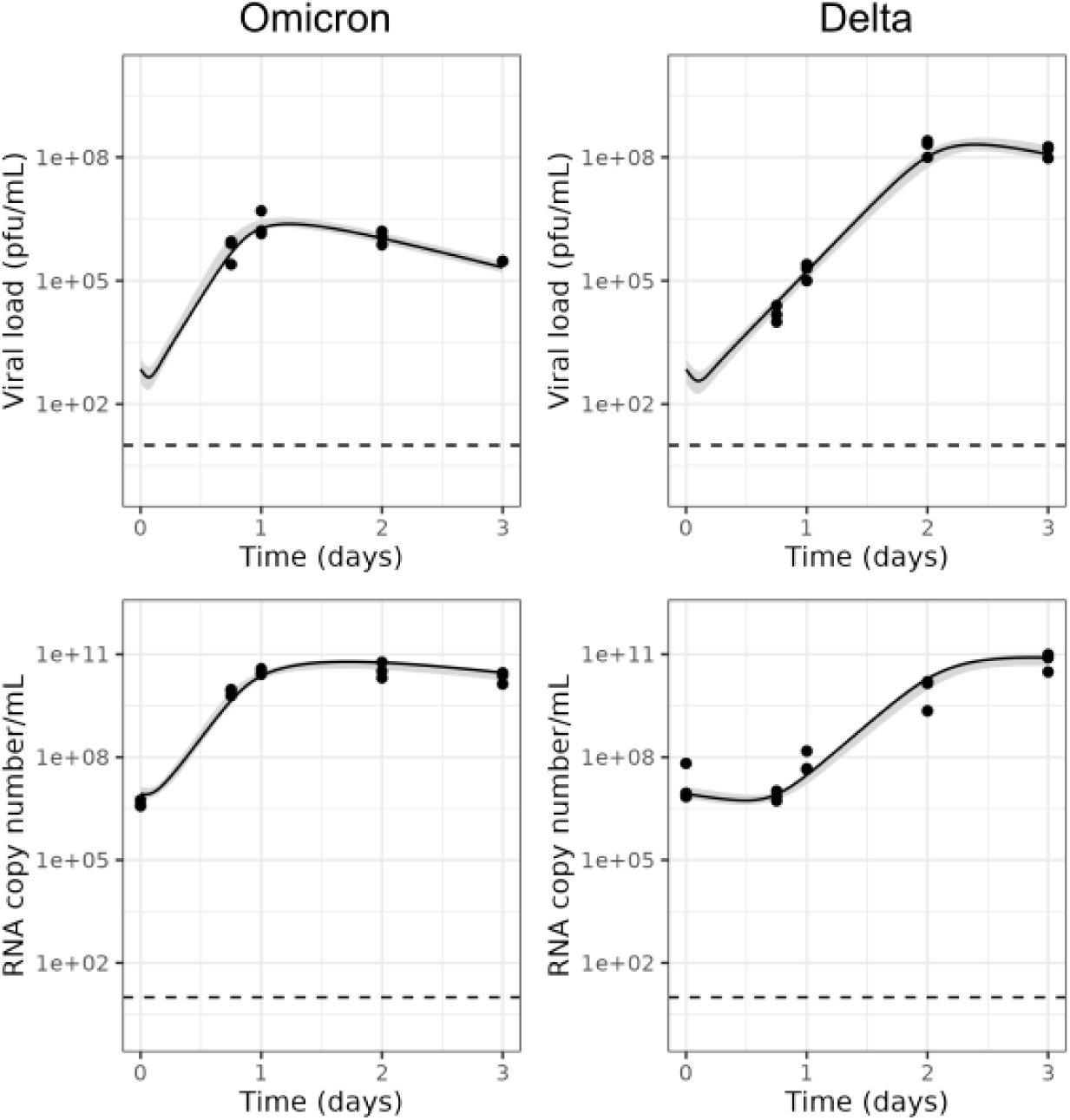
Model 1 fits to data from hNECs without drugs. Dots show the data, black lines show the maximum likelihood fit, shaded areas show the 95% credible interval (CrI) and dotted lines show the limit of detection.

**Figure 6:**
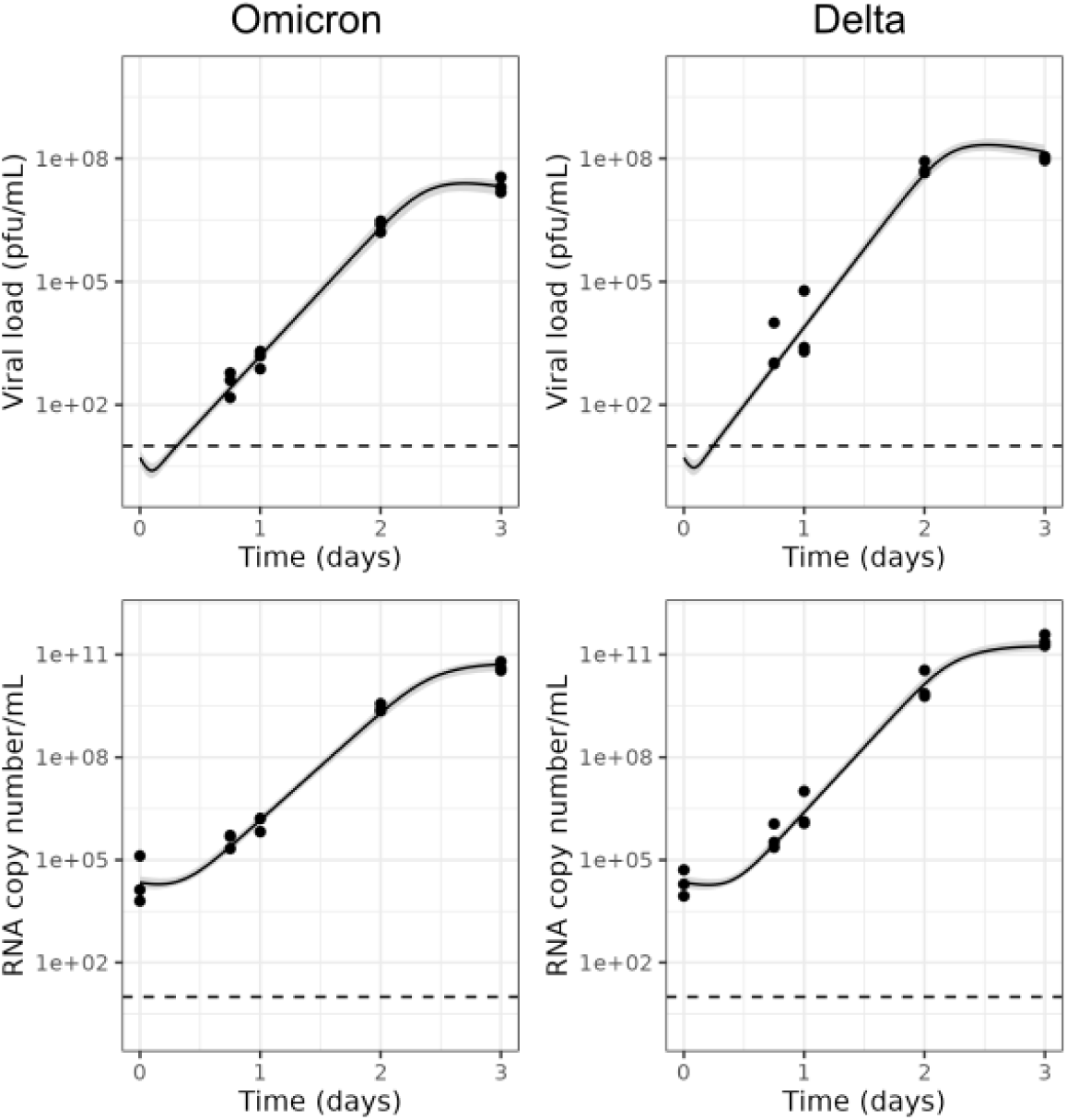
Model 1 fits to data from Calu-3 cells without drugs. Dots show the data, black lines show the maximum likelihood fit, shaded areas show the 95% credible interval (CrI) and dotted lines show the limit of detection.

For Calu-3 cells, we estimated a higher target cell infection rate for Omicron than Delta (9.23e-07 (95% CrI: 6.17e-07, 1.36e-06) cell (pfu/mL)^-1^ day^-1^ for Omicron, 1.42e-07 (95% CrI: 9.80e-08, 2.05e-07) cell (pfu/mL)^-1^ day^-1^ for Delta) and lower infectious virus production rate for Omicron than Delta (4.13e+02 (95% CrI: 2.85e+02,6.07e+02) cell (pfu/mL)^-1^ day^-1^ for Omicron, 3.73e+03 (95% CrI: 2.67e+03,5.32e+03) cell (pfu/mL)^-1^ day^-1^ for Delta). In contrast with hNECs, our results indicate an overall increase in viral fitness of Delta relative to Omicron in Calu-3 cells as we estimated a higher basic reproduction number of Delta (36.67 (95% CrI: 33.69, 39.81)) compared to Omicron (24.23 (95% CrI: 22.14, 26.70)) in this cell type. In addition, we estimated a larger growth rate of Delta relative to Omicron in Calu-3 cells, with a growth rate of 8.78 day^-1^ (95% CrI: 8.40, 9.15) for Delta compared with 7.25 day^-1^ (95% CrI: 6.86, 7.60) for Omicron (for details see Supplementary Table 2).

For both cell types, the target cell infection rate (*β*), which is positively correlated with the probability of a virion successfully infecting a cell, was higher for Omicron than Delta. The infectious virus production rate (ω_Inf_), which is directly proportional to the burst size (the total number of infectious virions produced by one infected cell), was higher for Delta than Omicron. However, the magnitude of each of these differences meant that the basic reproduction number and growth rate, which are both functions of the target cell infection rate and the infectious virus production rate (see Supplementary Methods), were higher for Omicron in hNECs, but higher for Delta in Calu-3 cells (Supplementary Table 2).

#### Model 2

When fitting Model 2 to the data, we found the same qualitative result that Omicron’s *R*_0_ is higher than Delta’s in hNECs, but lower in Calu-3 cells (Figures 7 and 8). In hNECs, *R*_0_ for Omicron was estimated to be 106.59 (95% CrI: 85.92, 134.72), while for Delta it was estimated to be 21.61 (95% CrI: 18.97, 25.30). In Calu-3 cells, *R*_0_ for Omicron was estimated to be 16.53 (95% CrI: 15.15, 17.99), and for Delta was estimated to be 27.87 (95% CrI: 25.60, 30.34). *R*_0_ estimates were comparable between Models 1 and 2, except for Omicron in hNECs, where the *R*_0_ estimates obtained with Model 2 were higher. This discrepancy occurred because Model 2 predicts that Omicron can use both TMPRSS2-dependent and TMPRSS2-independent pathways in hNECs (see next section), whereas for all other strain and cell combinations only TMPRSS2-dependent pathways are active. When only TMPRSS2-dependent pathways are active, then in the absence of drug, Models 1 and Model 2 are the same. All parameter estimates are presented in Supplementary Table 1.

**Figure 7.**
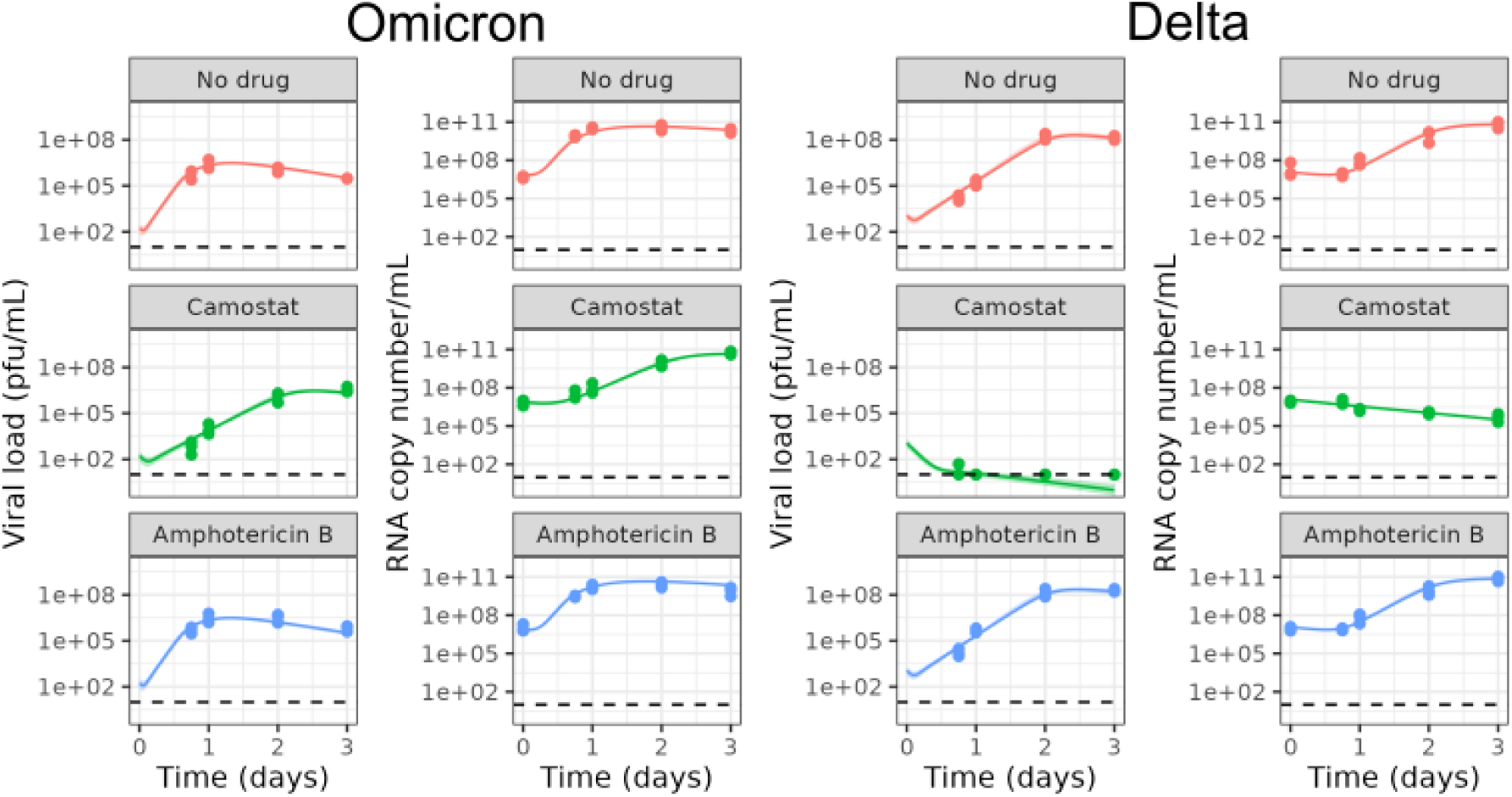
Model 2 fits to data from hNECs with and without drugs. Dots show the data, black lines show the maximum likelihood fit, shaded areas show the 95% credible interval (CrI) and dotted lines show the limit of detection. Markers on the lines represent measurements below the limit of detection.

**Figure 8.**
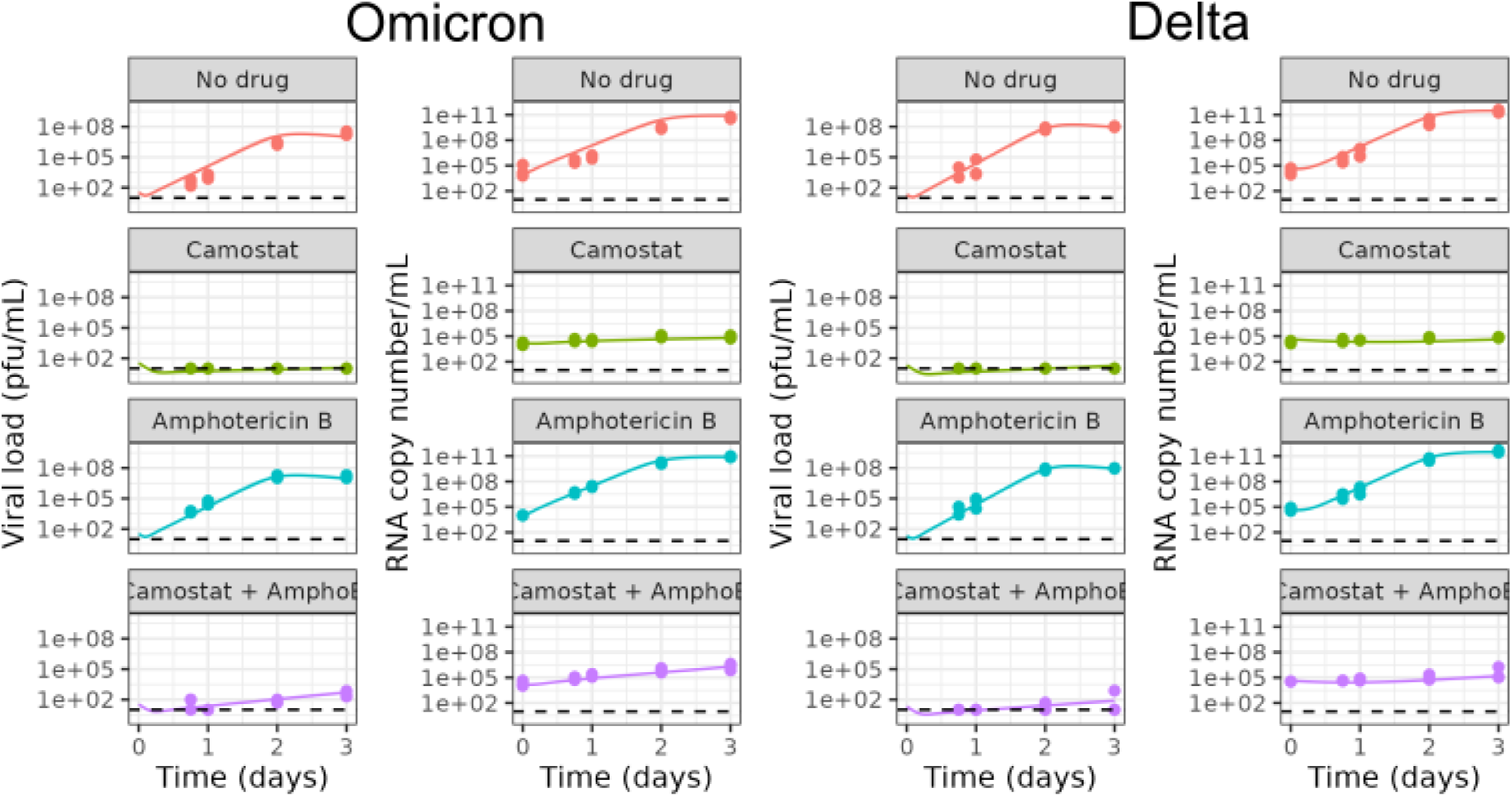
Model 2 fits to data from Calu-3 cells with and without drugs. Dots show the data, black lines show the maximum likelihood fit, shaded areas show the 95% credible interval (CrI) and dotted lines show the limit of detection. Markers on the lines represent measurements below the limit of detection.

### Role of Cell Entry Pathways

As Camostat inhibits serine proteases including TMPRSS2, Camostat-sensitive viruses must be able to use TMPRSS2-independent pathways for cell entry; thus, the fit of Model 2 to the data in the presence of Camostat provides information on the role of cell entry pathways. We found that in hNECs, Omicron was able to utilise both TMPRSS2-dependent and TMPRSS2-independent pathways. The median estimates of *R*_0*T*_ and *R*_0*E*_ were 92.13 (95% CrI: 72.95, 118.27) and 15.44 (95% CrI 12.86, 18.52) respectively (Supplementary Table 3). On the other hand, in hNECs, Delta was unable to effectively use TMPRSS2-independent pathways, as *R*_0*E*_ < 1 (median 0.15, 95% CrI: (0.07, 0.30)). In Calu-3s, neither virus was able to use TMPRRS2-independent pathways effectively. *R*_0*E*_ was estimated to be slightly greater than 1 for both Omicron (median 1.34 (95% CrI: 1.14, 1.55)) and Delta (median 1.71 (95% CrI: 1.47, 1.98)) but these small values of *R*_0_ are insufficient for the viral load to exceed the limit of detection during the time course of the experiment (Figure 8 Camostat panels). Note that for Omicron in hNECs, the estimate for *R*_0*T*_ is comparable to the overall *R*_0_ for Model 1. This is because the increase in viral load due to TMPRSS2-dependent pathways is much faster than through TMPRSS2-independent pathway, so the TMPRSS2-dependent pathway is the main contributor to the initial exponential increase, from which Model 1 estimates *R*_0_.

It has been suggested that Omicron’s use of TMPRRS2-independent pathways explains its fitness advantage over Delta^12,16,17^. However, even the TMPRSS2-specific basic reproduction number is higher for Omicron than Delta in hNECs, with a median *R*_0*T*_ equal to 92.13 (95% CrI: 72.95, 118.27) for Omicron compared to 21.46 (95% CrI:18.89, 25.05) for Delta. In fact, *R*_0*T*_ alone for Omicron is greater than the overall *R*_0_ for Delta. Therefore, it is primarily Omicron’s more efficient use of TMPRSS2-dependent pathways, rather than its utilisation of TMPRSS2-independent pathways, which gives it a fitness advantage over Delta in hNECs.

### Role of endosomal restriction

As Amphotericin B inhibits restrictions to virus entry imposed by endosomal restriction factors, the fit of Model 2 to the Amphotericin B data provides information on the role of endosomal restriction factors in shaping the viral dynamics observed. The parameters *f_E_* and *f_T_* capture the degree of inhibition of TMPRSS2-independent and TMPRSS2-dependent pathways by endosomal restriction factors respectively, and Amphotericin B is assumed to set these parameters to 0. Comparing across cell types, inhibition by endosomal restriction factors was higher in Calu-3 cells than hNECs across all strains and pathways (higher values of *f_E_* and *f_T_* for Calu-3 cells compared to hNEC cells in Supplementary Table 4), with the exception of TMPRSS2-independent pathways for Delta, where could not be estimated precisely. Inhibition was similar for both Omicron and Delta in hNECs, but Omicron was more inhibited by endosomal restriction factors than Delta in Calu-3 cells for both pathways (higher values of *f_E_* and *f_T_* for Omicron than Delta in Calu-3 cells in Supplementary Table 4). Comparing across pathways, *f_E_* and *f_T_* were similar for each virus strain in hNECs, but in Calu-3 cells,*f_E_* was higher than *f_T_* for both virus strains, suggesting more inhibition of TMPRSS2-independent pathways than of TMPRSS2-dependent pathways.

Given that TMPRSS2-independent pathways are susceptible to inhibition by endosomal restriction factors, and that previous variants were not able to use endosomal restriction pathways, one may hypothesise that Omicron has overcome endosomal restriction in hNECs, and that TMPRSS2-independent pathways would also be accessible by other virus-cell combinations if endosomal restriction were lifted. However, we predict that even if endosomal restriction is lifted for the three other virus-cell combinations (as per the Amphotericin B experiments) *R*_0*EX*_ remains low. This suggests that other factors contribute to Omicron’s increased use of TMPRSS2-independent pathways in hNECs. We predict that in the absence of endosomal restriction factors, Omicron would still have a higher overall basic reproduction number and growth rate than Delta in hNECs, but these quantities would be similar between variants in Calu-3 cells (*R*_0*X*_ in Supplementary Table 3 and *r_X_* in Supplementary Table 6).

### Changing the assumed eclipse phase duration

A caveat of the above results is that we have fixed the length of the eclipse phase. The eclipse phase in this model reflects the speed of viral replication with the cell. To date, studies which have measured the duration of the eclipse phase in the SARS-CoV-2 viral life cycle have used viruses which enter target cells through TMPRSS2-dependent pathways only^41,42^. Thus, the duration of the eclipse phase for TMPRSS2-independent pathways is unclear and our knowledge of this component of the viral life cycle is limited. To address this, we conducted a sensitivity analysis around the duration of the eclipse phase. In Model 1, *τ* was set to 6, 4, or 2 day^-1^. In Model 2, *τ_T_* was set to 6, 4 or 2 day^-1^, and *τ_E_* was set to 4, 2 or 1 day^-1^. We found that for Model 1, regardless of the duration of the eclipse phase assumed – as long as it was the same between strains – Omicron had a higher *R*_0_ than Delta in hNECs, and Delta had a higher *R*_0_ than Omicron in Calu-3 cells (Supplementary Table 4). Also, all estimated *R*_0_ values decreased as the duration of the eclipse phase decreased, as expected theoretically. This is because if, at the start of infection, cells start producing virus sooner due to a short eclipse phase, then each individual infected cell needs to lead to fewer secondary infections to reproduce the dynamics observed. Similarly, for Model 2, we found that regardless of the values of *τ_E_* and *τ_T_*, as long as these were assumed to be the same between strains, Omicron had a higher *R*_0_ than Delta in hNECs, and Delta had a higher *R*_0_ than Omicron in Calu-3 cells; only Omicron in hNECs used TMPRSS2-independent pathways efficiently; and endosomal restriction only had a significant effect in Calu-3 cells (Supplementary Table 5). However, allowing the duration of the eclipse phase to be different between strains, no longer allows us to draw conclusions about whether *R*_0_ is greater for Omicron or Delta (Supplementary Tables 3 and 5). On the other hand, estimates of the growth rate *r* only varied slightly with the assumed duration of the eclipse phase for both models (Supplementary Tables 4 and 6).

## Discussion

By fitting a simple within-host model to viral kinetics data for Omicron and Delta in hNECs and Calu-3 cells we found that Omicron has a fitness advantage over Delta in hNECs, but Delta has a fitness advantage over Omicron in Calu-3 cells, as measured by both the growth rate and basic reproduction number of the virus. These findings are consistent with previously published studies showing faster replication of Omicron compared with Delta in human nasal epithelial cells^14,16^, and faster replication of Delta compared with Omicron in the ex vivo explant cultures of human lungs^12^ and Calu-3 cells^43^.

We estimated that Omicron had a higher rate of infection *β* in both cell types, which could be linked to increased ACE2 binding affinity, likely due to mutations in the spike protein. Evidence is mixed as to whether Omicron has a higher ACE2 binding affinity than Delta. Cameroni et al.^44^ found that Omicron has a higher human ACE2 binding affinity than Delta, but Mannar et al.^45^ found that ACE2 binding affinity is similar for Omicron and Delta, though higher than the ancestral strain. On the other hand, Zhang et al.^46^ found that the Omicron spike protein required a higher level of ACE2 than Delta for efficient membrane fusion. We also estimated that Omicron had a lower rate of infectious virus production *ω_Inf_* than Delta for both cell types. Changes to the rate of viral production could be due to mutations in either spike or non-spike proteins. The basic reproduction number is positively correlated with both the rate of infection *β* and the rate of infectious virus production *ω_Inf_* in our models. In hNECs, the increase in the rate of infection *β* for Omicron was greater than the decrease in infectious virus production *ω_Inf_*, resulting in a larger basic reproduction number and growth rate for Omicron compared to Delta; the converse was true in Calu-3 cells.

Previous studies have proposed that TMPRSS2-independent pathways which are only available to Omicron are responsible for its faster growth in hNECs. Our study found that Omicron can use TMPRSS2-independent pathways in hNECs, consistent with the previous studies. However, the growth rate and basic reproduction number for TMPRSS2-independent pathways was low and insufficient to explain the overall fitness advantage for Omicron over Delta in hNECs. On the other hand, the growth rate and basic reproduction number for TMPRSS2-dependent pathways was also higher for Omicron than Delta in hNECs and was sufficient to explain the overall fitness advantage. Evidence for the continued importance of TMPRSS2-dependent pathways for Omicron was provided by a study by Metzdorf et al.^47^ which found that the growth of Omicron in the nose and lung was attenuated in TMPRSS2 knockout mice. In Calu-3 cells, neither Omicron nor Delta can use TMPRSS2-independent pathways, thereby providing further insight into how these viruses use different pathways to enter target cells.

We estimated the degree of inhibition in viral growth imposed by endosomal restriction factors by fitting our model to data where the cells were treated with Amphotericin B, as Amphotericin B inhibits the action of endosomal restriction factors. The main hypotheses tested were i) whether Omicron was able to evade endosomal restriction in hNECs; ii) whether this evasion enabled Omicron to use TMPRSS2-independent pathways; and iii) whether this was responsible for Omicron’s growth advantage. We found that the fitness of Delta in hNECs without endosomal restriction, as measured by the basic reproduction number *R*_0*X*_ and the growth rate *r_X_*, was still lower than that of Omicron in hNECs with endosomal restriction. Hence, reduced endosomal restriction alone do not explain Omicron’s fitness advantage in hNECs. Although we were able to quantify the degree of inhibition by endosomal restriction factors for both pathways for Omicron in hNECs, we were unable to do so for Delta, as doing so requires either the infectious viral load to be above the limit of detection in the presence of Camostat (as was the case for Omicron in hNECs), or requires data on viral growth in the presence of both drugs (as was the case in Calu-3 cells). Hence, we were unable to make a direct comparison at a pathway-specific level between Omicron and Delta in hNECs. In Calu-3 cells, we found that Omicron was more inhibited by endosomal restriction factors than Delta, for either TMPRSS2-dependent or TMPRSS2-independent pathways.

The model has several limitations. First, we fixed the infected cell death rate, the rate at which viruses lose infectivity, the RNA degradation rate and the duration of eclipse phases, according to values from the literature. It is currently unclear whether the duration of the eclipse phase differs between variants. To test this, we conducted a sensitivity analysis where the model was fitted assuming different lengths of the eclipse phase. This sensitivity analysis showed that if eclipse phase lengths were assumed to be the same between Omicron and Delta, while the estimated values of *R*_0_ changed with the assumed length of the eclipse phase, the trends and relationships in *R*_0_ and the growth rate estimates *r* between Omicron and Delta (for example, that Omicron has a higher *R*_0_ and growth rate estimate *r* than Delta in hNECs) remained unchanged. Allowing the eclipse phase to be different between strains, no longer allows us to draw conclusions about whether *R*_0_ is greater for Omicron or Delta (Supplementary Tables 3 and 5), but the differences in *r* are robust. This result reflects the finding that if the eclipse phase is allowed to vary, the growth rate *r* is identifiable but *R*_0_ is not. Future studies could measure the duration of the eclipse phase for each cell type and virus strain using single-cycle growth kinetics experiments.

We also did not test the effect of changing the assumed values of the infected cell or virus decay rates. Studies have shown that Omicron survives for longer on surfaces than the ancestral strain^48^ and Delta^49^, so it is plausible that extracellular virus may also differ in stability in our experiments. Moreover, fixing the parameters using values from previous studies may underestimate the uncertainty in *R*_0_ and *r*.

Another limitation of the model is that it assumes 100% effectiveness for both drugs. Assuming 100% effectiveness of Camostat means interpreting all remaining viral growth as due to TMPRSS2-independent pathways, so if the drug is less than 100% effective, the contribution of TMPRSS2-independent pathways will be overestimated. Assuming 100% effectiveness of Amphotericin B means interpreting viral growth in the presence of Amphotericin B as the viral growth in the complete absence of endosomal restriction. If the drug were less than 100% effective, the viral load in the complete absence of endosomal restriction could be higher than that suggested by the data, so the effect of endosomal restriction would be underestimated.

Last, the structure of the model is such that the initial growth rate is independent of the inoculum size. It is plausible that differences in growth rate between hNECs and Calu-3 cells observed in the experiments were partly due to the differing inoculum sizes used, as a higher inoculum could, for example, trigger endosomal restriction factors more quickly – an effect which was not modelled. A higher multiplicity of infection (MOI) was used for hNECs as a lower MOI did not always successfully infect these primary cells; repeating the Calu-3 experiments at the same higher MOI would strengthen confidence in our comparison between cell types.

The models developed in this study can inform future studies using more complex models. Models capturing more immune components and how the virus spreads within the host are useful for understanding contributors to disease severity and the effects of antivirals^32–34^, but because they have many parameters, it is often difficult to determine their values by fitting them to data. This study can help parameterize these models.

The models developed in this study can be applied to different SARS-CoV-2 virus strains to help enhance our understanding of transmission potential of new strains as the virus continues to evolve. While between-strain differences in the speed of viral replication is typically assessed by comparing viral titre measurements at individual timepoints, adopting a mathematical modelling approach using longitudinal timepoints enables us to estimate key characteristics of different strains such as the growth rate and viral fitness, thus providing a deeper insight into the factors shaping observed differences in viral dynamics at the individual and population level.

## Acknowledgements

CMC and ID acknowledge funding from the MRC Centre for Global Infectious Disease Analysis (reference MR/R015600/1), jointly funded by the UK Medical Research Council (MRC) and the UK Foreign, Commonwealth & Development Office (FCDO), under the MRC/FCDO Concordat agreement and is also part of the EDCTP2 programme supported by the European Union. AWCY acknowledges funding from an Imperial College Research Fellowship. ID acknowledges funding by the Wellcome Trust and Royal Society (grant number 213494/Z/18/Z). This study was conducted as part of G2P-UK National Virology consortium funded by MRC/UKRI (MR/W005611/1). For the purpose of open access, the authors have applied a CC BY public copyright licence to any Author Accepted Manuscript version arising from this submission.

## Author Contributions

CMC, AWCY, WSB and ID conceived the study; JCB and KS generated the *in-vitro* data; CMC, AWCY and ID conceived the mathematical models; CMC performed the analysis for Model 1; AWCY performed the analysis for Model 2; CMC, AWCY, JCB, TPP, WSB, and ID contributed to the interpretation of the results; CMC and AWCY wrote the first draft of the manuscript; all authors reviewed and approved the final version of the manuscript.

## Data Availability

All data is available at https://github.com/ada-w-yan/deltaomicronmodelling

## Code Availability

All code is available at https://github.com/ada-w-yan/deltaomicronmodelling

