## Supplementary Methods and Tables for "Modelling the viral dynamics of the SARS-CoV-2 Delta and Omicron variants in different cell types"

#### Contents

**1. Supplementary Methods** .......... 2
**2. Supplementary Tables** .......... 6

### 1. Supplementary Methods

**Model 1**

The dynamics of Model 1 are described by the following set of deterministic equations:

$$\frac{dT}{dt} = -\beta \mathrm{V}_{\mathrm{Inf}} T$$

$$\frac{dL}{dt} = \beta \mathrm{V}_{\mathrm{Inf}} T - \tau\, L$$

$$\frac{dI}{dt} = \tau\, L - \delta I$$

$$\frac{d\mathrm{V}_{\mathrm{Inf}}}{dt} = \omega_{\mathrm{Inf}} I - \kappa_{\mathrm{Inf}} \mathrm{V}_{\mathrm{Inf}} - \beta \mathrm{V}_{\mathrm{Inf}} T$$

$$\frac{d\mathrm{V}_{\mathrm{RNA}}}{dt} = \omega_{\mathrm{RNA}} I - \kappa_{\mathrm{RNA}} \mathrm{V}_{\mathrm{RNA}} - \beta V_{Inf} T$$

Parameter definitions are in Table 1. The basic reproduction number ($R_0$) for this model is defined as the mean number of infected cells produced by each infected cell over its lifespan at under disease-free conditions (at the start of infection, time t*) and is given by:

$$R_0 = \frac{\beta \mathrm{T}^* \omega_{\mathrm{Inf}}}{\delta(\kappa_{\mathrm{Inf}} + \beta T^*)}$$

where $T^*$ is the initial number of target cells. The initial growth rate (*r*) is obtained by calculating the largest eigenvalue of the Jacobian matrix obtained when linearising the equations above around the disease-free equilibrium.

**Model 2**

The dynamics of Model 2 are described by the following set of equations:

$$\frac{dT_-}{dt} = -\beta_E(1 - f_E) T_- V_{Inf}$$

$$\frac{dT_+}{dt} = -[\beta_E(1 - f_E) + \beta_T(1 - f_T)] T_+ V_{Inf}$$

$$\frac{dL_E}{dt} = \beta_E(1 - f_E)(T_- + T_+) V_{Inf} - \tau_E L_E$$

$$\frac{dL_T}{dt} = \beta_T(1 - f_T) T_+ V_{Inf} - \tau_T L_T$$

$$\frac{dI}{dt} = \tau_E L_E + \tau_T L_T - \delta I$$

$$\frac{dV_{Inf}}{dt} = \omega_{Inf} I - \left[\kappa_{Inf} + \beta_E(1 - f_E)(T_- + T_+) + \beta_T(1 - f_T)T_+\right]V_{Inf}$$

$$\frac{dV_{RNA}}{dt} = \omega_{RNA} I - \kappa_{RNA} V_{RNA} - [\beta_E(1 - f_E)(T_- + T_+) + \beta_T(1 - f_T)T_+]V_{Inf}$$

Parameter definitions are in Table 1. The basic reproduction number for this model is

$$R_0 = \frac{[\beta_E(1 - f_E)T_-^* + \beta_E(1 - f_E)T_+^* + \beta_T(1 - f_T)T_+^*]\omega_{Inf}}{\delta[\kappa_{Inf} + \beta_E(1 - f_E)T_-^* + \beta_E(1 - f_E)T_+^* + \beta_T(1 - f_T)T_+^*]}$$

We also define pathway-specific $R_0$s. $R_{0E}$ is the number of infectious virions produced by one infectious virus, if only the TMPRSS2-independent pathway were active; it is computed by setting $\beta_T$= 0 in the above definition of $R_0$. Similarly, $R_{0T}$ is the number of infectious virions produced by one infectious virus, if only the TMPRSS2-dependent pathway were active; it is computed by setting $\beta_E$= 0 in the above definition of $R_0$. Pathway-specific growth rates are determined similarly.

In addition, we can define both pathway-specific and overall $R_0$ in the absence of IFITM, denoted $R_{0EX}$, $R_{0TX}$ and $R_{0X}$ by setting $f_E = f_T = 0$. For example,

$$R_{0EX} = \frac{[\beta_E T_-^* + \beta_E T_+^*]\omega_{Inf}}{\delta[\kappa_{Inf} + \beta_E T_-^* + \beta_E T_+^*]}$$

In both models, we use the scaling factor $\mathrm{s_{Inf}}$ to quantify the amount of viral inoculum (measured in pfu) that successfully results in infecting target cells by setting:

$$\mathrm{V_{Inf}}(0) = \mathrm{s_{Inf}} \mathrm{V_{Inf}^*}$$

where $\mathrm{V_{Inf}^*}$ denotes the viral inoculum. We use the scaling factor $\mathrm{s_{RNA}}$ to determine the corresponding amount of non-infectious virus present at the beginning of each experiment by setting:

$$\mathrm{V_{RNA}}(0) = \mathrm{s_{RNA}} \mathrm{V_{Inf}^*}$$

**Likelihood and Inferential Framework**

For each strain, the model was simultaneously fitted to both plaque-assay (infectious) and qPCR (non-infectious) virus measurements collected in individual wells using the Metropolis-Hastings algorithm[1].

All parameters except for $f_E$ and $f_T$ were estimated on a $\log_{10}$ scale using uniform prior distributions and univariate Normal proposal distribution. Details of fixed parameter values and the prior distributions assumed for the estimated model parameters are provided in Table 1. The 95% Credible

Intervals (CrI) around the central estimates were generated using 1,000 parameter sets sampled from the posterior distribution.

| Symbol | Parameter Description | Parameter Value | Reference | Prior |
|---|---|---|---|---|
| $1/\tau$ | Duration of eclipse phase (days) | 0.25 | [2] | n/a |
| $\beta, \beta_E, \beta_T$ | Target cell infection rate (cells per pfu/mL day$^{-1}$) | Estimated | n/a | log10(x) ~ U[-12, -2] |
| $\omega_{Inf}$ | Infectious virus production rate (pfu/mL per infectious cell, day$^{-1}$) | Estimated | n/a | log10(x) ~ U[0, 10] |
| $\omega_{RNA}$ | Total virus production rate (RNA copy number/mL per infectious cell, day$^{-1}$) | Estimated | n/a | log10(x) ~ U[0, 10] |
| $f_E, f_T$ | Decrease in infectivity due to endosomal restriction factors | Estimated | n/a | x ~ U[0, 1] |
| $\delta$ | Infected cell decay rate (day$^{-1}$) | 1.7 | [2] | n/a |
| $\kappa_{Inf}$ | Infectious virus decay rate (day$^{-1}$) | 10 | [2] | n/a |
| $\kappa_{RNA}$ | Non-infectious virus decay rate (day$^{-1}$) | 1.2 | [3] | n/a |
| $V^*_{Inf}$ | Initial infectious virus inoculum (pfu/mL) | 1.2e3 (Calu-3), 2.5e4 (hNEC) | n/a | n/a |
| $s_{Inf}$ | Scaling factor for infectious virus inoculum | Estimated | n/a | log10(x) ~ U[-20, 1] |
| $s_{RNA}$ | Scaling factor for non-infectious virus inoculum (RNA copy number per (pfu/mL)) | Estimated | n/a | log10(x) ~ U[-20, 4] |
| T(0) | Initial number of target cells | 1.2e6 (Calu-3), 1.3e4 (hNEC) | [4] | n/a |
| $T_+(0)$ | Initial number of TMRPSS2+ cells ($T(0) = T_+(0) + T_-(0)$) | 1.2e6 (Calu-3), 4672 (hNEC) | [4] | n/a |

**Supplementary Table 1: Model 1 Parameters**

As cells of the same type received the same quantity of inoculum in experiments using both the Delta and Omicron variants, $s_{Inf}$ and $s_{RNA}$ were constrained to be the same for Omicron and Delta for a given cell type for Model 1. However, this constraint hindered convergence for Model 2 and thus was not used for this model. All other parameters were treated as variant-specific.

For calculation of the likelihood, let $\widehat{V}_{Inf_{ijk}}$ and $\widehat{V}_{RNA_{ijk}}$ denote the observed $\log_{10}$ plaque-assay and qPCR viral measurements for variant *i* in well *j* on day *k*, with the corresponding model predicted values represented by $V_{Inf_{ijk}}$ and $V_{RNA_{ijk}}$. We assumed that the $\log_{10}$ measurements were distributed according to a Normal distribution with variance $\sigma^2$, and measurements below the limit of detection (LOD) were accounted for in the likelihood using the cumulative distribution function, with the likelihood given by:

$$\mathrm{L} = \prod_{\mathrm{Y}=\{V_{Inf},V_{RNA}\}} \prod_{i=1}^{2}\prod_{j=1}^{J}\prod_{k=1}^{K} \phi\left(\left(\mathrm{Y}_{ijk} \mid \hat{\mathrm{Y}}_{ijk}, \sigma^2\right)\right)^{c_{\mathrm{Y}ijk}} \, \Phi\left(\left(LOD_{\mathrm{Y}} \mid \hat{\mathrm{Y}}_{ijk}, \sigma^2\right)\right)^{1-c_{\mathrm{Y}ijk}}$$

where $\phi$ and $\Phi$ denote the probability density function (pdf) and cumulative distribution function (cdf) of the Normal distribution respectively, $Y$ denotes the variable of interest, the indicator function $c_{\mathrm{Y}ijk}$ is 1 if the observed value was above the limit of detection $\left(Y_{ijk} > LOD_Y\right)$ and 0 otherwise, $J$ denotes the total number of wells and $K$ denotes the total number of measurements. The limit of detection was 10 pfu/mL for infectious virus and 10 RNA copy number/mL for viral RNA.

We fix $\sigma$ to the standard deviation of

$$\log_{10} V_{s,d,t,m} - \frac{\sum_{n=1}^{N} \log_{10} V_{s,t,d,n}}{N}$$

across all strains, drugs, timepoints and replicates, where $s$ is the strain, $d$ is the drug (no drug, Camostat or Amphotericin B), $t$ is the timepoint, $m$ is the replicate number (1, 2 or 3), and N = 3 is the total number of replicates. Where a value of $V_{s,d,t,m}$ is below the limit of detection, that combination of strain, drug and timepoint is excluded from the calculation for all $m$. The calculation was done separately for infectious virus and RNA copy number, leading to values of the standard deviation of 0.24 and 0.22.

### 2. Supplementary Tables

**Supplementary Table 2. Median and 95% CrI (within parentheses) for the estimated parameters.**

| | **Omicron hNEC** | |
|---|---|---|
| | **Model 1** | **Model 2** |
| $\beta$ | 2.13e-05 (1.54e-05, 3.07e-05) | NA |
| $\beta_E$ | NA | 4.45e-06 (3.29e-06, 6.48e-06) |
| $\beta_T$ | NA | 7.09e-05 (5.17e-05, 1.03e-04) |
| $\omega_{Inf}$ | 4.07e+03 (3.13e+03, 5.50e+03) | 5.09e+03 (4.18e+03, 6.25e+03) |
| $\omega_{RNA}$ | 1.72e+07 (1.24e+07, 2.28e+07) | 1.46e+07 (1.20e+07, 1.80e+07) |
| $f_E$ | NA | 9.62e-02 (4.15e-03, 3.35e-01) |
| $f_T$ | NA | 3.22e-02 (9.50e-04, 1.52e-01) |
| $s_{Inf}$ | 2.73e-02 (1.31e-02, 5.22e-02) | 5.98e-03 (2.78e-03, 1.27e-02) |
| $s_{RNA}$ | 3.63e+02 (2.45e+02, 5.44e+02) | 2.82e+02 (2.02e+02, 3.93e+02) |
| $R_0$ | 65.04 (51.94, 82.71) | 106.59 (85.92, 134.72) |
| r | 11.61 (10.43, 12.99) | 14.56 (13.20, 16.17) |
| | **Delta hNEC** | |
| | **Model 1** | **Model 2** |
| $\beta$ | 8.89e-08 (5.91e-08, 1.28e-07) | NA |
| $\beta_E$ | NA | 4.51e-10 (1.10e-10, 2.75e-09) |
| $\beta_T$ | NA | 8.51e-08 (5.53e-08, 1.30e-07) |
| $\omega_{Inf}$ | 3.42e+05 (2.41e+05, 5.04e+05) | 9.74e+05 (6.89e+05, 1.41e+06) |
| $\omega_{RNA}$ | 2.32e+07 (1.54e+07, 3.38e+07) | 5.76e+07 (3.95e+07, 8.15e+07) |
| $f_E$ | NA | 5.51e-01 (3.89e-02, 8.97e-01) |
| $f_T$ | NA | 3.86e-02 (2.07e-03, 1.18e-01) |
| $s_{Inf}$ | 2.73e-02 (1.31e-02, 5.22e-02) | 4.37e-02 (2.26e-02, 7.92e-02) |
| $s_{RNA}$ | 3.63e+02 (2.45e+02, 5.44e+02) | 4.66e+02 (3.78e+02, 5.69e+02) |
| $R_0$ | 23.13 (19.91, 27.23) | 21.61 (18.97, 25.30) |
| r | 6.87 (6.33, 7.50) | 6.62 (6.16, 7.21) |
| | **Omicron Calu-3** | |
| | **Model 1** | **Model 2** |
| $\beta$ | 9.23e-07 (6.17e-07, 1.36e-06) | NA |
| $\beta_E$ | NA | 1.27e-07 (9.15e-08, 1.74e-07) |
| $B_T$ | NA | 1.17e-06 (8.29e-07, 1.69e-06) |
| $\omega_{Inf}$ | 4.13e+02 (2.85e+02, 6.07e+02) | 3.46e+02 (2.62e+02, 4.64e+02) |
| $\omega_{RNA}$ | 1.82e+05 (1.19e+05, 2.69e+05) | 3.02e+05 (2.26e+05, 4.09e+05) |
| $f_E$ | NA | 5.62e-01 (5.07e-01, 6.15e-01) |
| $f_T$ | NA | 4.19e-01 (3.55e-01, 4.79e-01) |
| $s_{Inf}$ | 4.38e-03 (2.76e-03, 6.70e-03) | 2.81e-02 (1.95e-02, 4.06e-02) |

| | | |
|---|---|---|
| $s_{rna}$ | 1.88e+01 (1.27e+01, 2.77e+01) | 1.41e+01 (1.03e+01, 1.91e+01) |
| $R_0$ | 24.23 (22.14, 26.70) | 16.53 (15.15, 17.99) |
| r | 7.22 (6.86, 7.60) | 5.79 (5.50, 6.10) |
| | **Delta Calu-3** | |
| | **Model 1** | **Model 2** |
| β | 1.42e-07 (9.80e-08, 2.05e-07) | NA |
| $\beta_E$ | NA | 1.31e-08 (9.41e-09, 1.71e-08) |
| $\beta_T$ | NA | 1.84e-07 (1.34e-07, 2.40e-07) |
| $\omega_{Inf}$ | 3.73e+03 (2.67e+03, 5.32e+03) | 2.56e+03 (1.99e+03, 3.45e+03) |
| $\omega_{RNA}$ | 6.37e+05 (4.46e+05, 9.46e+05) | 1.08e+06 (8.20e+05, 1.46e+06) |
| $f_E$ | NA | 2.72e-01 (1.77e-01, 3.58e-01) |
| $f_T$ | NA | 1.99e-01 (1.30e-01, 2.65e-01) |
| $s_{Inf}$ | 4.38e-03 (2.76e-03, 6.70e-03) | 1.60e-02 (1.05e-02, 2.42e-02) |
| $s_{RNA}$ | 1.88e+01 (1.27e+01, 2.77e+01) | 3.60e+01 (2.70e+01, 4.68e+01) |
| $R_0$ | 36.67 (33.69, 39.81) | 27.87 (25.60, 30.34) |
| r | 8.78 (8.40, 9.15) | 7.62 (7.29, 7.97) |

**Supplementary Table 3: $R_0$ estimates (median and 95% CrI within parentheses) through different pathways in Model 2 for Omicron and Delta.**

| | hNEC | | Calu-3 | |
|---|---|---|---|---|
| | Omicron | Delta | Omicron | Delta |
| $R_{0T}$ | 92.13 (72.95, 118.27) | 21.46 (18.89, 25.05) | 15.38 (14.09, 16.75) | 26.21 (24.10, 28.52) |
| $R_{0E}$ | 15.44 (12.86, 18.52) | 0.15 (0.07, 0.30) | 1.34 (1.14, 1.55) | 1.71 (1.47, 1.98) |
| $R_0$ | 106.59 (85.92, 134.72) | 21.61 (18.97, 25.30) | 16.53 (15.15, 17.99) | 27.87 (25.60, 30.34) |
| $R_{0TX}$ | 96.34 (76.07, 124.47) | 22.45 (19.63, 26.47) | 25.06 (22.59, 28.00) | 32.52 (29.74, 36.24) |
| $R_{0EX}$ | 17.25 (13.83, 23.66) | 0.34 (0.10, 1.65) | 3.04 (2.71, 3.38) | 2.36 (2.05, 2.67) |
| $R_{0X}$ | 112.89 (91.22, 142.58) | 22.96 (19.87, 27.47) | 27.40 (24.86, 30.39) | 34.77 (31.78, 38.67) |

**Supplementary Table 4. Median and 95% CI for all fitted parameter values for Model 1 with different lengths of the eclipse phase.**

| | **Omicron hNEC** | | |
|---|---|---|---|
| | **$\tau$=2 day$^{-1}$** | **$\tau$=4 day$^{-1}$** | **$\tau$=6 day$^{-1}$** |
| $\beta$ | 5.52E-05(3.62E-05,9.54E-05) | 2.13E-05(1.54E-05,3.07E-05) | 1.46E-05(1.05E-05,2.10E-05) |
| $\omega_{Inf}$ | 3.56E+03(2.66E+03,4.71E+03) | 4.07E+03(3.13E+03,5.50E+03) | 4.22E+03(3.16E+03,5.64E+03) |
| $\omega_{RNA}$ | 1.62E+07(1.20E+07,2.18E+07) | 1.72E+07(1.24E+07,2.28E+07) | 1.72E+07(1.29E+07,2.40E+07) |
| $s_{Inf}$ | 2.22E-02(8.70E-03,4.94E-02) | 2.73E-02(1.31E-02,5.22E-02) | 2.79E-02(1.29E-02,5.65E-02) |
| $s_{RNA}$ | 3.63E+02(2.47E+02,5.21E+02) | 3.63E+02(2.45E+02,5.44E+02) | 3.60E+02(2.45E+02,5.40E+02) |
| *R0* | 139.86(101.26,205.00) | 65.04(51.94,82.71) | 46.68(36.18,58.23) |
| *r* | 12.86(11.09,15.34) | 11.61(10.43,12.99) | 11.32(9.96,12.62) |

| | **Delta hNEC** | | |
|---|---|---|---|
| | **$\tau$=2 day$^{-1}$** | **$\tau$=4 day$^{-1}$** | **$\tau$=6 day$^{-1}$** |
| $\beta$ | 1.49E-07(9.44E-08,2.37E-07) | 8.89E-08(5.91E-08,1.28E-07) | 6.60E-08(4.49E-08,9.68E-08) |
| $\omega_{Inf}$ | 3.56E+05(2.44E+05,5.35E+05) | 3.42E+05(2.41E+05,5.04E+05) | 3.52E+05(2.43E+05,4.99E+05) |
| $\omega_{RNA}$ | 2.47E+07(1.63E+07,3.73E+07) | 2.32E+07(1.54E+07,3.38E+07) | 2.37E+07(1.59E+07,3.46E+07) |
| $s_{Inf}$ | 2.22E-02(8.70E-03,4.94E-02) | 2.73E-02(1.31E-02,5.22E-02) | 2.79E-02(1.29E-02,5.65E-02) |
| $s_{RNA}$ | 3.63E+02(2.47E+02,5.21E+02) | 3.63E+02(2.45E+02,5.44E+02) | 3.60E+02(2.45E+02,5.40E+02) |
| *R0* | 40.49(33.28,51.61) | 23.13(19.91,27.23) | 17.84(15.46,21.10) |
| *r* | 7.12(6.45,8.02) | 6.87(6.33,7.50) | 6.78(6.24,7.46) |

| | **Omicron Calu-3** | | |
|---|---|---|---|
| | **$\tau$=2 day$^{-1}$** | **$\tau$=4 day$^{-1}$** | **$\tau$=6 day$^{-1}$** |
| $\beta$ | 1.37E-06(9.41E-07,2.13E-06) | 9.23E-07(6.17E-07,1.36E-06) | 7.11E-07(5.05E-07,1.06E-06) |
| $\omega_{Inf}$ | 4.78E+02(3.23E+02,6.67E+02) | 4.13E+02(2.85E+02,6.07E+02) | 4.15E+02(2.86E+02,5.79E+02) |
| $\omega_{RNA}$ | 2.06E+05(1.34E+05,3.19E+05) | 1.82E+05(1.19E+05,2.69E+05) | 1.79E+05(1.18E+05,2.62E+05) |
| $s_{Inf}$ | 4.50E-03(2.93E-03,7.09E-03) | 4.38E-03(2.76E-03,6.70E-03) | 4.26E-03(2.66E-03,6.55E-03) |
| $s_{RNA}$ | 1.85E+01(1.25E+01,2.77E+01) | 1.88E+01(1.27E+01,2.77E+01) | 1.86E+01(1.21E+01,2.84E+01) |
| *R0* | 39.45(35.97,43.51) | 24.23(22.14,26.70) | 19.15(17.62,21.00) |
| *r* | 7.25(6.90,7.61) | 7.22(6.86,7.60) | 7.20(6.87,7.59) |

| | Delta Calu-3 | | |
|---|---|---|---|
| | **$\tau$=2 day$^{-1}$** | **$\tau$=4 day$^{-1}$** | **$\tau$=6 day$^{-1}$** |
| $\beta$ | 2.26E-07(1.54E-07,3.31E-07) | 1.42E-07(9.80E-08,2.05E-07) | 1.08E-07(7.54E-08,1.59E-07) |
| $\omega_{Inf}$ | 4.02E+03(2.75E+03,5.82E+03) | 3.73E+03(2.67E+03,5.32E+03) | 3.74E+03(2.60E+03,5.45E+03) |
| $\omega_{RNA}$ | 7.08E+05(4.60E+05,1.04E+06) | 6.37E+05(4.46E+05,9.46E+05) | 6.39E+05(4.38E+05,9.43E+05) |
| $s_{Inf}$ | 4.50E-03(2.93E-03,7.09E-03) | 4.38E-03(2.76E-03,6.70E-03) | 4.26E-03(2.66E-03,6.55E-03) |
| $s_{RNA}$ | 1.85E+01(1.25E+01,2.77E+01) | 1.88E+01(1.27E+01,2.77E+01) | 1.86E+01(1.21E+01,2.84E+01) |
| *R0* | 62.14(56.08,68.23) | 36.67(33.69,39.81) | 28.19(25.95,30.58) |
| *r* | 8.82(8.39,9.22) | 8.77(8.40,9.15) | 8.75(8.38,9.14) |

**Supplementary Table 5. Median and 95% CI for $R_0$ for Model 2 with different lengths of the eclipse phase.**

| $R_0$ | | | | | |
|---|---|---|---|---|---|
| $\tau_E$ | $\tau_T$ | hNEC Omicron | Calu-3 Omicron | hNEC Delta | Calu-3 Delta |
| 1 | 2 | 340.79 (244.39, 523.09) | 26.23 (23.84, 28.78) | 36.85 (31.17, 46.42) | 47.37 (42.70, 52.42) |
| 1 | 4 | 174.81 (131.11, 255.91) | 17.06 (15.53, 18.58) | 21.20 (18.55, 24.75) | 29.68 (26.81, 32.84) |
| 1 | 6 | 133.45 (104.09, 194.46) | 13.97 (12.78, 15.33) | 16.62 (14.81, 18.97) | 23.60 (21.61, 25.86) |
| 2 | 2 | 106.59 (85.92, 134.72) | 16.53 (15.15, 17.99) | 21.61 (18.97, 25.30) | 27.87 (25.60, 30.34) |
| 2 | 4 | 128.04 (102.08, 165.98) | 16.70 (15.32, 18.17) | 21.46 (18.67, 25.16) | 28.64 (25.93, 31.37) |
| 2 | 6 | 96.71 (77.58, 122.09) | 13.59 (12.54, 14.85) | 16.67 (14.80, 19.17) | 22.59 (20.76, 24.53) |
| 4 | 2 | 224.24 (173.37, 302.04) | 25.95 (23.77, 28.33) | 37.56 (31.57, 50.52) | 45.42 (41.29, 50.10) |
| 4 | 4 | 106.59 (85.92, 134.72) | 16.53 (15.15, 17.99) | 21.61 (18.97, 25.30) | 27.87 (25.60, 30.34) |
| 4 | 6 | 78.14 (64.20, 95.91) | 13.36 (12.35, 14.49) | 16.80 (14.93, 19.41) | 21.97 (20.24, 23.81) |

| $R_{0X}$ | | | | | |
|---|---|---|---|---|---|
| $\tau_E$ | $\tau_T$ | hNEC Omicron | Calu-3 Omicron | hNEC Delta | Calu-3 Delta |
| 1 | 2 | 381.25 (265.61, 596.43) | 45.92 (41.29, 51.59) | 39.64 (33.24, 51.40) | 61.36 (54.73, 69.53) |
| 1 | 4 | 195.52 (145.08, 300.54) | 29.61 (26.87, 32.68) | 22.49 (19.50, 27.46) | 37.73 (34.03, 42.87) |
| 1 | 6 | 149.52 (113.05, 229.85) | 24.16 (22.02, 26.61) | 17.64 (15.49, 21.26) | 29.73 (27.00, 33.14) |
| 2 | 2 | 112.89 (91.22, 142.58) | 27.40 (24.86, 30.39) | 22.96 (19.87, 27.47) | 34.77 (31.78, 38.67) |
| 2 | 4 | 138.21 (109.33, 178.86) | 28.14 (25.52, 31.17) | 22.84 (19.69, 27.31) | 35.99 (32.52, 40.09) |
| 2 | 6 | 104.59 (83.59, 133.89) | 22.47 (20.63, 24.92) | 17.63 (15.51, 20.65) | 28.09 (25.68, 30.98) |
| 4 | 2 | 239.85 (186.41, 325.68) | 44.53 (40.02, 50.13) | 40.16 (33.21, 55.45) | 58.61 (52.26, 65.79) |
| 4 | 4 | 112.89 (91.22, 142.58) | 27.40 (24.86, 30.39) | 22.96 (19.87, 27.47) | 34.77 (31.78, 38.67) |
| 4 | 6 | 82.48 (68.03, 100.76) | 21.57 (19.73, 23.79) | 17.72 (15.62, 20.85) | 27.08 (24.85, 29.90) |

| $R_{0E}$ | | | | | |
|---|---|---|---|---|---|
| $\tau_E$ | $\tau_T$ | hNEC Omicron | Calu-3 Omicron | hNEC Delta | Calu-3 Delta |
| 1 | 2 | 58.54 (41.88, 97.76) | 1.96 (1.56, 2.42) | 0.21 (0.10, 0.46) | 2.83 (2.26, 3.50) |
| 1 | 4 | 56.73 (41.04, 92.19) | 1.96 (1.56, 2.40) | 0.17 (0.08, 0.36) | 2.87 (2.26, 3.55) |
| 1 | 6 | 54.83 (40.92, 88.55) | 1.95 (1.54, 2.41) | 0.18 (0.09, 0.34) | 2.86 (2.31, 3.50) |
| 2 | 2 | 15.44 (12.86, 18.52) | 1.34 (1.14, 1.55) | 0.15 (0.07, 0.30) | 1.71 (1.47, 1.98) |
| 2 | 4 | 26.48 (21.36, 34.61) | 1.53 (1.26, 1.82) | 0.15 (0.07, 0.32) | 2.07 (1.72, 2.46) |
| 2 | 6 | 26.73 (21.52, 35.11) | 1.50 (1.26, 1.81) | 0.15 (0.07, 0.29) | 2.07 (1.73, 2.47) |
| 4 | 2 | 15.64 (12.95, 19.41) | 1.35 (1.15, 1.57) | 0.17 (0.07, 0.46) | 1.71 (1.46, 2.00) |
| 4 | 4 | 15.44 (12.86, 18.52) | 1.34 (1.14, 1.55) | 0.15 (0.07, 0.30) | 1.71 (1.47, 1.98) |
| 4 | 6 | 15.37 (12.93, 18.88) | 1.32 (1.13, 1.55) | 0.14 (0.07, 0.30) | 1.71 (1.47, 1.98) |

| $R_{0EX}$ | | | | | |
|---|---|---|---|---|---|
| $\tau_E$ | $\tau_T$ | hNEC Omicron | Calu-3 Omicron | hNEC Delta | Calu-3 Delta |
| 1 | 2 | 74.67 (46.47, 167.27) | 5.80 (5.01, 6.70) | 0.56 (0.14, 4.35) | 4.34 (3.56, 5.19) |
| 1 | 4 | 69.44 (46.23, 147.72) | 5.80 (5.05, 6.60) | 0.38 (0.12, 2.80) | 4.39 (3.62, 5.32) |
| 1 | 6 | 66.81 (45.39, 128.17) | 5.80 (5.02, 6.73) | 0.40 (0.12, 2.86) | 4.35 (3.66, 5.19) |
| 2 | 2 | 17.25 (13.83, 23.66) | 3.04 (2.71, 3.38) | 0.34 (0.10, 1.65) | 2.36 (2.05, 2.67) |
| 2 | 4 | 31.10 (22.99, 46.81) | 3.93 (3.45, 4.44) | 0.36 (0.11, 1.88) | 3.02 (2.54, 3.54) |
| 2 | 6 | 30.97 (23.40, 49.16) | 3.93 (3.47, 4.46) | 0.35 (0.10, 1.82) | 2.99 (2.57, 3.50) |
| 4 | 2 | 17.88 (14.06, 25.86) | 3.05 (2.73, 3.43) | 0.38 (0.10, 1.78) | 2.35 (2.05, 2.70) |
| 4 | 4 | 17.25 (13.83, 23.66) | 3.04 (2.71, 3.38) | 0.34 (0.10, 1.65) | 2.36 (2.05, 2.67) |
| 4 | 6 | 17.08 (13.85, 22.84) | 3.01 (2.71, 3.35) | 0.29 (0.10, 1.32) | 2.35 (2.05, 2.68) |

| $R_{0T}$ | | | | | |
|---|---|---|---|---|---|
| $\tau_E$ | $\tau_T$ | hNEC Omicron | Calu-3 Omicron | hNEC Delta | Calu-3 Delta |
| 1 | 2 | 122.46 (89.53, 181.43) | 15.36 (14.04, 16.71) | 21.03 (18.45, 24.42) | 26.91 (24.49, 29.54) |
| 1 | 4 | 81.39 (62.34, 116.44) | 12.25 (11.25, 13.36) | 16.44 (14.69, 18.69) | 20.82 (19.17, 22.66) |

| | | | | | |
|---|---|---|---|---|---|
| 1 | 6 | 92.13 (72.95, 118.27) | 15.38 (14.09, 16.75) | 21.46 (18.89, 25.05) | 26.21 (24.10, 28.52) |
| 2 | 2 | 103.41 (80.52, 137.08) | 15.40 (14.12, 16.72) | 21.31 (18.57, 24.88) | 26.63 (24.17, 29.13) |
| 2 | 4 | 71.09 (55.78, 91.37) | 12.26 (11.32, 13.39) | 16.53 (14.70, 18.88) | 20.58 (18.95, 22.30) |
| 2 | 6 | 210.47 (160.73, 287.04) | 24.90 (22.76, 27.19) | 37.40 (31.48, 50.07) | 43.80 (39.83, 48.29) |
| 4 | 2 | 92.13 (72.95, 118.27) | 15.38 (14.09, 16.75) | 21.46 (18.89, 25.05) | 26.21 (24.10, 28.52) |
| 4 | 4 | 63.27 (51.16, 79.28) | 12.19 (11.24, 13.22) | 16.65 (14.84, 19.16) | 20.30 (18.75, 21.98) |
| 4 | 6 | 122.46 (89.53, 181.43) | 15.36 (14.04, 16.71) | 21.03 (18.45, 24.42) | 26.91 (24.49, 29.54) |

| $R_{0TX}$ | | | | | |
|---|---|---|---|---|---|
| $\tau_E$ | $\tau_T$ | hNEC Omicron | Calu-3 Omicron | hNEC Delta | Calu-3 Delta |
| 1 | 2 | 316.28 (214.61, 513.65) | 41.98 (37.58, 47.47) | 38.75 (32.62, 49.46) | 57.31 (51.22, 65.06) |
| 1 | 4 | 129.87 (93.95, 194.92) | 25.00 (22.63, 27.90) | 21.94 (19.14, 25.95) | 33.51 (30.29, 37.97) |
| 1 | 6 | 85.54 (65.24, 123.67) | 19.31 (17.64, 21.38) | 17.08 (15.17, 19.49) | 25.51 (23.26, 28.38) |
| 2 | 2 | 96.34 (76.07, 124.47) | 25.06 (22.59, 28.00) | 22.45 (19.63, 26.47) | 32.52 (29.74, 36.24) |
| 2 | 4 | 108.94 (84.52, 146.60) | 25.08 (22.59, 27.95) | 22.33 (19.37, 26.34) | 33.07 (29.93, 36.86) |
| 2 | 6 | 74.32 (57.58, 96.02) | 19.25 (17.54, 21.47) | 17.16 (15.24, 19.69) | 25.18 (23.07, 27.78) |
| 4 | 2 | 223.93 (170.10, 310.35) | 42.56 (38.01, 48.12) | 39.67 (32.90, 54.13) | 56.40 (50.25, 63.41) |
| 4 | 4 | 96.34 (76.07, 124.47) | 25.06 (22.59, 28.00) | 22.45 (19.63, 26.47) | 32.52 (29.74, 36.24) |
| 4 | 6 | 65.95 (52.86, 82.54) | 19.08 (17.38, 21.24) | 17.34 (15.33, 20.10) | 24.81 (22.73, 27.45) |

**Supplementary Table 6. Median and 95% CI for r for Model 2 with different lengths of the eclipse phase.**

| $r$ | | | | | |
|---|---|---|---|---|---|
| $\tau_E$ | $\tau_T$ | hNEC Omicron | Calu-3 Omicron | hNEC Delta | Calu-3 Delta |
| 1 | 2 | 18.40 (15.79, 22.50) | 5.73 (5.43, 6.03) | 6.78 (6.23, 7.60) | 7.64 (7.26, 8.03) |
| 1 | 4 | 16.31 (14.24, 19.32) | 5.66 (5.35, 5.94) | 6.53 (6.07, 7.10) | 7.62 (7.24, 8.01) |
| 1 | 6 | 15.63 (13.89, 18.36) | 5.61 (5.31, 5.92) | 6.48 (6.06, 6.98) | 7.57 (7.22, 7.94) |
| 2 | 2 | 14.56 (13.20, 16.17) | 5.79 (5.50, 6.10) | 6.62 (6.16, 7.21) | 7.62 (7.29, 7.97) |
| 2 | 4 | 15.19 (13.64, 17.11) | 5.72 (5.42, 6.01) | 6.59 (6.10, 7.18) | 7.62 (7.23, 7.99) |
| 2 | 6 | 14.77 (13.28, 16.49) | 5.67 (5.38, 5.99) | 6.50 (6.06, 7.04) | 7.57 (7.22, 7.91) |
| 4 | 2 | 16.10 (14.43, 18.29) | 5.88 (5.59, 6.17) | 6.87 (6.28, 7.96) | 7.67 (7.31, 8.05) |
| 4 | 4 | 14.56 (13.20, 16.17) | 5.79 (5.50, 6.10) | 6.62 (6.16, 7.21) | 7.62 (7.29, 7.97) |
| 4 | 6 | 14.13 (12.86, 15.56) | 5.73 (5.44, 6.02) | 6.54 (6.10, 7.11) | 7.58 (7.25, 7.93) |

| $r_X$ | | | | | |
|---|---|---|---|---|---|
| $\tau_E$ | $\tau_T$ | hNEC Omicron | Calu-3 Omicron | hNEC Delta | Calu-3 Delta |
| 1 | 2 | 19.20 (16.31, 23.74) | 7.73 (7.29, 8.26) | 7.01 (6.42, 7.94) | 8.65 (8.19, 9.19) |
| 1 | 4 | 16.90 (14.74, 20.21) | 7.58 (7.18, 8.05) | 6.71 (6.22, 7.37) | 8.57 (8.13, 9.13) |
| 1 | 6 | 16.13 (14.34, 19.04) | 7.49 (7.11, 7.93) | 6.65 (6.19, 7.21) | 8.49 (8.08, 8.98) |
| 2 | 2 | 14.94 (13.56, 16.57) | 7.77 (7.35, 8.25) | 6.85 (6.32, 7.54) | 8.56 (8.16, 9.03) |
| 2 | 4 | 15.66 (14.05, 17.65) | 7.67 (7.23, 8.14) | 6.79 (6.27, 7.45) | 8.56 (8.13, 9.04) |
| 2 | 6 | 15.18 (13.63, 16.97) | 7.56 (7.17, 8.04) | 6.69 (6.22, 7.27) | 8.48 (8.09, 8.94) |
| 4 | 2 | 16.62 (14.94, 18.96) | 7.96 (7.50, 8.51) | 7.11 (6.46, 8.36) | 8.70 (8.23, 9.20) |
| 4 | 4 | 14.94 (13.56, 16.57) | 7.77 (7.35, 8.25) | 6.85 (6.32, 7.54) | 8.56 (8.16, 9.03) |
| 4 | 6 | 14.47 (13.19, 15.89) | 7.63 (7.24, 8.08) | 6.74 (6.27, 7.37) | 8.49 (8.11, 8.95) |

| $r_E$ | | | | | |
|---|---|---|---|---|---|
| $\tau_E$ | $\tau_T$ | hNEC Omicron | Calu-3 Omicron | hNEC Delta | Calu-3 Delta |
| 1 | 2 | 6.46 (5.46, 8.27) | 0.47 (0.29, 0.65) | -0.64 (-0.80, -0.38) | 0.79 (0.59, 1.00) |
| 1 | 4 | 6.36 (5.41, 8.05) | 0.47 (0.29, 0.64) | -0.69 (-0.83, -0.48) | 0.80 (0.59, 1.02) |
| 1 | 6 | 6.26 (5.40, 7.90) | 0.46 (0.28, 0.64) | -0.69 (-0.82, -0.50) | 0.80 (0.61, 1.00) |
| 2 | 2 | 5.47 (4.90, 6.08) | 0.33 (0.14, 0.52) | -1.28 (-1.49, -0.95) | 0.65 (0.45, 0.85) |
| 2 | 4 | 5.73 (5.10, 6.59) | 0.39 (0.20, 0.57) | -1.07 (-1.30, -0.76) | 0.72 (0.51, 0.92) |
| 2 | 6 | 5.76 (5.12, 6.64) | 0.38 (0.20, 0.57) | -1.08 (-1.31, -0.81) | 0.72 (0.52, 0.93) |
| 4 | 2 | 5.52 (4.92, 6.25) | 0.34 (0.16, 0.54) | -1.22 (-1.47, -0.69) | 0.65 (0.44, 0.87) |
| 4 | 4 | 5.47 (4.90, 6.08) | 0.33 (0.14, 0.52) | -1.28 (-1.49, -0.95) | 0.65 (0.45, 0.85) |
| 4 | 6 | 5.46 (4.92, 6.15) | 0.32 (0.14, 0.52) | -1.29 (-1.49, -0.96) | 0.65 (0.45, 0.85) |

| $r_{EX}$ | | | | | |
|---|---|---|---|---|---|
| $\tau_E$ | $\tau_T$ | hNEC Omicron | Calu-3 Omicron | hNEC Delta | Calu-3 Delta |
| 1 | 2 | 7.28 (5.76, 10.61) | 1.59 (1.41, 1.79) | -0.30 (-0.74, 1.24) | 1.24 (1.02, 1.45) |
| 1 | 4 | 7.03 (5.74, 10.01) | 1.59 (1.42, 1.77) | -0.45 (-0.77, 0.78) | 1.25 (1.04, 1.48) |
| 1 | 6 | 6.89 (5.69, 9.37) | 1.59 (1.41, 1.79) | -0.43 (-0.76, 0.80) | 1.24 (1.05, 1.45) |
| 2 | 2 | 5.84 (5.12, 6.97) | 1.54 (1.35, 1.73) | -0.88 (-1.39, 0.60) | 1.12 (0.91, 1.32) |
| 2 | 4 | 6.24 (5.31, 7.67) | 1.56 (1.37, 1.74) | -0.70 (-1.19, 0.61) | 1.18 (0.96, 1.40) |

| | | | | | |
|---|---|---|---|---|---|
| 2 | 6 | 6.22 (5.36, 7.86) | 1.56 (1.38, 1.75) | -0.71 (-1.22, 0.57) | 1.17 (0.98, 1.39) |
| 4 | 2 | 5.96 (5.18, 7.31) | 1.54 (1.36, 1.75) | -0.81 (-1.38, 0.71) | 1.12 (0.91, 1.33) |
| 4 | 4 | 5.84 (5.12, 6.97) | 1.54 (1.35, 1.73) | -0.88 (-1.39, 0.60) | 1.12 (0.91, 1.32) |
| 4 | 6 | 5.81 (5.13, 6.84) | 1.52 (1.35, 1.71) | -0.97 (-1.40, 0.32) | 1.11 (0.91, 1.32) |

| $r_T$ | | | | | |
|---|---|---|---|---|---|
| $\tau_E$ | $\tau_T$ | hNEC Omicron | Calu-3 Omicron | hNEC Delta | Calu-3 Delta |
| 1 | 2 | 17.75 (15.10, 21.93) | 5.62 (5.33, 5.92) | 6.77 (6.22, 7.58) | 7.51 (7.14, 7.90) |
| 1 | 4 | 15.50 (13.44, 18.47) | 5.53 (5.24, 5.82) | 6.52 (6.06, 7.08) | 7.48 (7.11, 7.86) |
| 1 | 6 | 14.75 (13.02, 17.38) | 5.48 (5.18, 5.79) | 6.47 (6.05, 6.97) | 7.43 (7.09, 7.79) |
| 2 | 2 | 13.62 (12.23, 15.26) | 5.55 (5.25, 5.85) | 6.60 (6.14, 7.17) | 7.38 (7.05, 7.72) |
| 2 | 4 | 14.36 (12.81, 16.31) | 5.54 (5.25, 5.84) | 6.57 (6.09, 7.15) | 7.44 (7.06, 7.81) |
| 2 | 6 | 13.85 (12.34, 15.56) | 5.49 (5.20, 5.80) | 6.49 (6.05, 7.01) | 7.38 (7.04, 7.72) |
| 4 | 2 | 15.34 (13.63, 17.61) | 5.66 (5.37, 5.95) | 6.84 (6.26, 7.90) | 7.44 (7.09, 7.81) |
| 4 | 4 | 13.62 (12.23, 15.26) | 5.55 (5.25, 5.85) | 6.60 (6.14, 7.17) | 7.38 (7.05, 7.72) |
| 4 | 6 | 13.10 (11.83, 14.56) | 5.47 (5.18, 5.76) | 6.52 (6.09, 7.07) | 7.32 (7.00, 7.65) |

| $r_{TX}$ | | | | | |
|---|---|---|---|---|---|
| $\tau_E$ | $\tau_T$ | hNEC Omicron | Calu-3 Omicron | hNEC Delta | Calu-3 Delta |
| 1 | 2 | 18.37 (15.47, 22.97) | 7.54 (7.09, 8.08) | 6.96 (6.38, 7.86) | 8.49 (8.04, 9.04) |
| 1 | 4 | 15.92 (13.76, 19.09) | 7.36 (6.95, 7.84) | 6.68 (6.19, 7.31) | 8.39 (7.97, 8.95) |
| 1 | 6 | 15.09 (13.29, 17.86) | 7.25 (6.86, 7.72) | 6.61 (6.17, 7.13) | 8.31 (7.90, 8.80) |
| 2 | 2 | 13.90 (12.47, 15.61) | 7.38 (6.96, 7.87) | 6.76 (6.28, 7.39) | 8.27 (7.89, 8.74) |
| 2 | 4 | 14.70 (13.10, 16.82) | 7.38 (6.94, 7.86) | 6.74 (6.23, 7.37) | 8.34 (7.91, 8.81) |
| 2 | 6 | 14.14 (12.53, 15.92) | 7.25 (6.85, 7.75) | 6.63 (6.18, 7.18) | 8.25 (7.87, 8.70) |
| 4 | 2 | 15.76 (13.96, 18.21) | 7.63 (7.15, 8.19) | 7.05 (6.41, 8.21) | 8.43 (7.97, 8.92) |
| 4 | 4 | 13.90 (12.47, 15.61) | 7.38 (6.96, 7.87) | 6.76 (6.28, 7.39) | 8.27 (7.89, 8.74) |
| 4 | 6 | 13.37 (12.01, 14.84) | 7.21 (6.82, 7.68) | 6.67 (6.21, 7.26) | 8.18 (7.80, 8.64) |
